# Map making: Constructing, combining, and inferring on abstract cognitive maps

**DOI:** 10.1101/810051

**Authors:** Seongmin A. Park, Douglas S. Miller, Hamed Nili, Charan Ranganath, Erie D. Boorman

**Author notes:** Lead contact: S. A. Park and E. D. Boorman.

## Abstract

Cognitive maps are thought to enable model-based inferences from limited experience that can guide novel decisions–a hallmark of behavioral flexibility. We tested whether the hippocampus (HC), entorhinal cortex (EC), and ventromedial prefrontal cortex (vmPFC)/medial orbitofrontal cortex (mOFC) organize abstract and discrete relational information into a cognitive map to guide novel inferences. Subjects learned the status of people in two separate unseen 2-D social hierarchies defined by competence and popularity piecemeal from binary comparisons, with each dimension learned on a separate day. Although only one dimension was ever behaviorally relevant, multivariate activity patterns in HC, EC and vmPFC/mOFC were linearly related to the Euclidean distance between people in the mentally reconstructed 2-D space. Hubs created unique comparisons between the two hierarchies, enabling inferences between novel pairs of people. We found that both behavior and neural activity in EC and vmPFC/mOFC reflected the Euclidean distance to the retrieved hub, which was reinstated in HC. These findings reveal how abstract and discrete relational structures are represented, combined, and enable novel inferences in the human brain.

## INTRODUCTION

To form rich world models, sparse observations often sampled from separate experiences need to be integrated into a coherent representation. There has been a recent surge of interest in the long-standing theory that the hippocampus (HC) and entorhinal cortex (EC) may organize spatial and non-spatial relational information into such a ‘cognitive map’ for goal-directed behavior (Behrens et al., 2018; Bellmund, Gärdenfors, Moser, & Doeller, 2018; Cohen, 2015; Constantinescu, O’Reilly, & Behrens, 2016; Howard Eichenbaum & Cohen, 2014; Ekstrom & Ranganath, 2018; Hafting, Fyhn, Molden, Moser, & Moser, 2005; Moser, Kropff, & Moser, 2008; O’Keefe & Nadel, 1978; Schiller et al., 2015; Schuck, Cai, Wilson, & Niv, 2016; Tolman, 1948; Wikenheiser & Schoenbaum, 2016). While past studies have identified neural signals in the HC and EC indicative of a cognitive map primarily using continuous task dimensions with online sensory feedback during task performance (e.g. visual, auditory, vestibular) (Aronov, Nevers, & Tank, 2017; Bao et al., 2019; Constantinescu et al., 2016; Doeller, Barry, & Burgess, 2010; Howard Eichenbaum & Cohen, 2014; Hafting et al., 2005; Nau, Navarro Schröder, Bellmund, & Doeller, 2018; O’Keefe & Nadel, 1978; Theves, Fernandez, & Doeller, 2019), many important everyday decisions involve discrete entities that vary along multiple abstract dimensions that are sampled piecemeal, one experience at a time, in the absence of continuous sensory feedback, such as with whom to collaborate or where to eat. Whether, and if so, how the brain constructs a cognitive map of abstract relationships between discrete entities from piecemeal experiences is unclear.

A powerful advantage of a cognitive map of an environment or task is the ability to make inferences from sparse observations that can dramatically accelerate learning and even guide novel decisions never faced before (Banino et al., 2018; Behrens et al., 2018; Jones et al., 2012; Stachenfeld, Botvinick, & Gershman, 2017; Tolman, 1948; Vikbladh et al., 2019), a hallmark of behavioral flexibility and a key challenge in artificial intelligence (Behrens et al., 2018; Kriete, Noelle, Cohen, & O’Reilly, 2013; Wang et al., 2018). This is in part because a cognitive map of a task space would in theory allows “shortcuts” and “novel routes” to be inferred, as in physical space. To provide a concrete example, understanding the structure of family trees allows one to infer new relationships, such as the following: because Sally is John’s sister and Sue is John’s daughter, Sue must be Sally’s niece without ever directly learning this relationship (**Fig. 1A**). Biologically inspired computational models show the map-like coding schemes found in the HC and EC can in principle enable agents to perform vector navigation, including planning new routes and finding shortcuts to a goal in physical space (Banino et al., 2018; Bush, Barry, Manson, & Burgess, 2015; Dordek, Soudry, Meir, & Derdikman, 2016; J. C. R. R. Whittington et al., 2018). In particular, so-called place cells in HC and grid cells in medial EC contain neural codes that permit calculation of predicted position (Moser et al., 2008; O’Keefe & Nadel, 1978; Stachenfeld et al., 2017), direction (Banino et al., 2018; Chadwick, Jolly, Amos, Hassabis, & Spiers, 2015), and Euclidean distance (Behrens et al., 2018; Bellmund et al., 2018; Howard et al., 2014) in physical space. Yet despite recent theoretical proposals (J. C. Whittington et al., 2019), empirical evidence concerning how neural representations of abstract cognitive maps relate to such direct novel inferences outside of physical space has been lacking.

**Figure 1.**
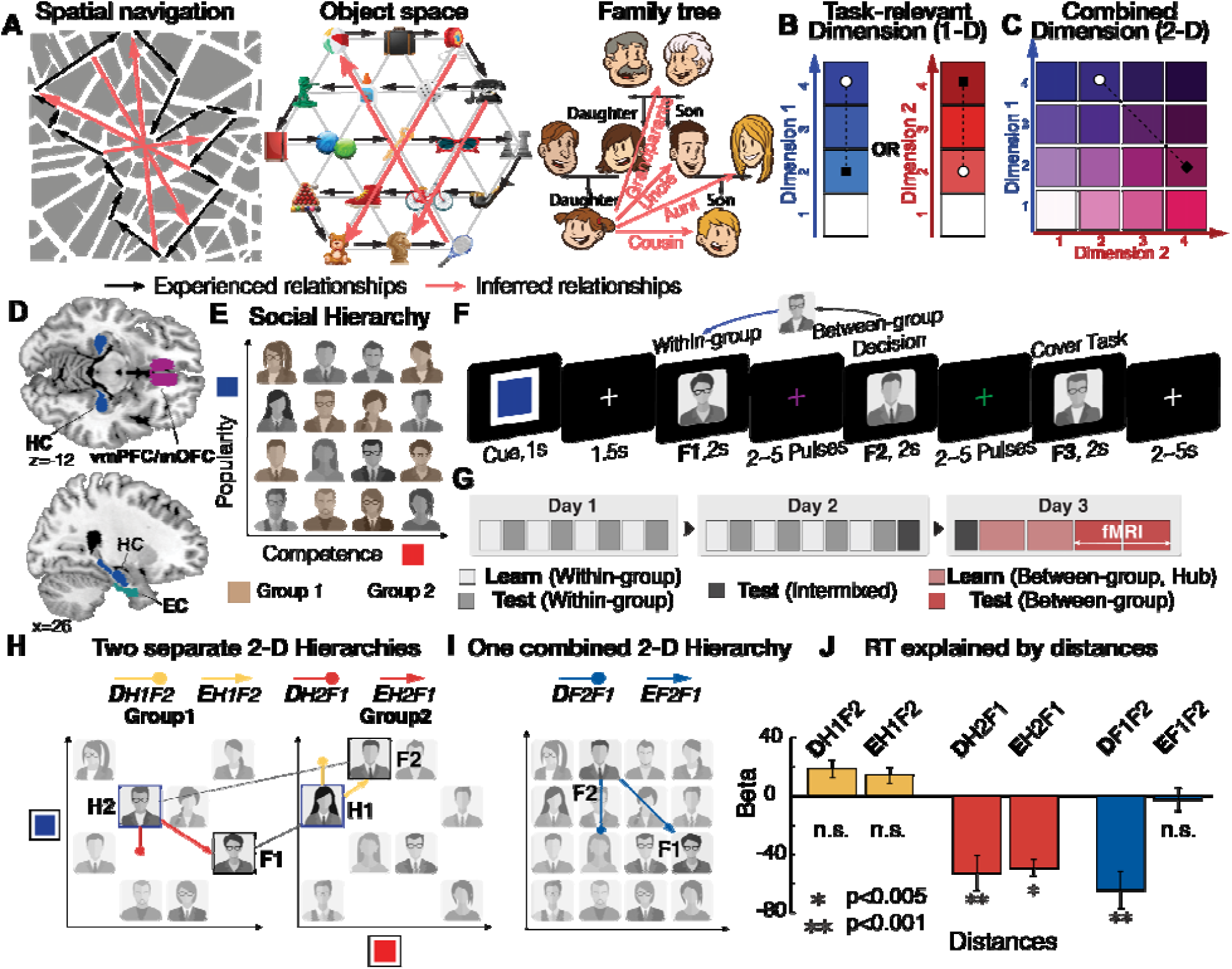
**A**. Examples of “zero-shot inferences” in physical space, transitions between objects, and family trees. Representing abstract relationships as a cognitive map allows making novel direct inferences that do not only rely on previously experienced associations. Black: experienced relationships; Red: inferred relationships. **B** and **C**. Two hypotheses concerning how the brain could represent and flexibly switch between different dimensions that characterize the same entities to guide inferences. **B**. The brain could construct two separate maps for representing the hierarchy of each dimension learned on a separate day and distinct regions could encode the one-dimensional (1-D) rank difference in the task-relevant dimension and the task-irrelevant dimension. **C**. Alternatively, the brain could construct a unified map consisting of two dimensions and encode the inferred Euclidean distance over the 2-D representation. **D**. Regions of interest (ROIs) generated independently from probabilistic maps in other studies (the vmPFC/mOFC (F.-X. Neubert et al., 2015), HC (Yushkevich et al., 2015), and EC (Amunts et al., 2005; Zilles & Amunts, 2010). All ROIs were defined bilaterally. **E**. Participants learned the rank of members of each of two groups (brown and gray) separately in two dimensions: competence and popularity. Crucially, subjects were never shown the 1- or 2-D structures but could infer them by making transitive inferences. **F**. Illustration of a trial of the fMRI experiment. Participants made inferences about the relative status of a novel pair (F1 and F2) in a given dimension (signaled by the Cue color). A cover task (to indicate the gender of the face stimulus, F3) followed at the end of every trial. **G**. On day 1 and day 2, during learning blocks participants learned within-group ranks of the two groups in each of two dimensions through binary decisions about the status of members who differed by only one rank level in a given dimension. Test blocks tested subjects’ knowledge of the two 1-D hierarchies that could be constructed using transitive inferences for each group separately. On day3, subjects learned from between-group comparisons limited to ‘hub’ individuals, which created a unique path between groups per person in each dimension. Subsequently, on day3, participants were asked to infer the unlearned between-group status while undergoing fMRI. **H**. Participants could use hubs to infer the relationship between novel pairs. Possible trajectories for two example inferences can be seen. Two distances are shown for each trajectory: the behaviorally-relevant 1-D distance (D) and the 2-D Euclidean distance (E). Subjects could use either of two trajectories: a forward inference from F1 to its hub (H1) that has a unique connection to F2 (1-D distance: D_H1F2_, Euclidean distance: E_H1F2_) which is shown in yellow; or a backward inference from F2 to its hub (H2) that has a unique connection to F1 (D_H2F1;_ E_H2F1_) which is shown in red. **I**. As alternative paths, subjects may not use the hubs, but instead compute the distance in the relevant dimension between F1 and F2 directly (D_F1F2_), or their Euclidean distance (E_F1F2_) in the combined cognitive map of two groups (blue). **J**. Multiple linear regression results show that both the rank distance (D_H2F1_) and the Euclidean distance from H2 (E_H2F1_), but not from H1, significantly explain variance in reaction times (RT), in addition to the direct distance between F1 and F2 (D_F1F2_), while competing with other distance terms.

A parallel literature based on recent studies focusing on the orbitofrontal cortex (OFC) has motivated a related theory that the OFC represents one’s current position in a cognitive map, not of physical space, but of task space (Schuck et al., 2016; Takahashi et al., 2017; Walton, Behrens, Buckley, Rudebeck, & Rushworth, 2010; Wikenheiser & Schoenbaum, 2016; Wilson, Takahashi, Schoenbaum, & Niv, 2014). Recent findings further suggest a specialized role for mOFC in representing all the latent (or perceptually unsignaled) components of the task space that define one’s current state in the task (Muller, Mars, Behrens, & O’Reilly, 2019; Schuck et al., 2016; Wilson et al., 2014). This function is proposed to play an important role both during learning (Takahashi et al., 2017; Walton et al., 2010) and choice (Jones et al., 2012; Stalnaker, Cooch, & Schoenbaum, 2015). Recent studies have indeed discovered that the OFC represents latent task states during learning and choice in support of this theory (Chan, Niv, & Norman, 2016; Schuck et al., 2016; Wikenheiser, Marrero-Garcia, & Schoenbaum, 2017), yet to our knowledge there has been little direct evidence of map-like representations (e.g. position, direction, or distance) of the task space in OFC. Moreover, whether this proposed OFC function would extend to representing a cognitive map of an abstract social space, or whether it would instead transfer to areas implicated in social cognition is unclear.

In addition to *representing* cognitive maps, both the HC and OFC have been implicated in model-based *inference*, such that distinct stimuli, or stimuli and rewards, that were not directly associated can be associated or integrated through an overlapping, shared associate (Jones et al., 2012; Koster et al., 2018; Kurth-Nelson, Economides, Dolan, & Dayan, 2016; Schlichting & Preston, 2014; Tompary & Davachi, 2017; Wimmer & Shohamy, 2012). Computational models have proposed how a related mechanism in the HC-EC system may additionally underlie transitive inferences about ordinal rank (Koster et al., 2018; Kumaran & McClelland, 2012) (though see (Frank, Rudy, Levy, & O’Reilly, 2005)). In addition to demonstrations that HC is necessary for transitive inferences (Howard Eichenbaum, Otto, & Cohen, 1996), studies in animal models have demonstrated that the OFC is necessary for model-based inferences based on previously learned associations, but not for decisions based on directly learned cached values (Jones et al., 2012). While these HC and OFC roles for associating or integrating individual items have been documented, how the brain constructs broader cognitive maps and makes direct inferences beyond chaining or integrating previously experienced elementary associations has been elusive. In particular, it is possible that similar mechanisms of integration, and/or distinct mechanisms that leverage an explicit representation of the relational structure of the space or task (J. C. Whittington et al., 2019) would allow for the integration of distinct relational structures into a single larger cognitive map from sparse observations (e.g. the integration of family trees through marriage) that even respects metric relationships (e.g. vector directions and distances) and enables direct inferences that have never been experienced before (e.g. Sue is Sally’s niece; **Fig. 1A**).

Here, we asked participants to learn two 2-D social hierarchies from the outcomes of binary decisions about individuals’ rank on either of the two dimensions, with each dimension learned on a different day. During fMRI participants were asked to make novel inferences about the relative status of a novel pair across hierarchies in only one dimension at a time. This manipulation meant subjects were required to flexibly switch between currently relevant and irrelevant social dimensions that described the same entities for decisions. We tested two non-mutually exclusive hypotheses concerning how the human brain could represent and flexibly switch between different dimensions that characterize the same entities to guide direct inferences. Based on previous decision-making studies showing behaviorally-relevant decision value signals in vmPFC/mOFC, and pending or currently irrelevant value signals in separate prefrontal areas (Boorman, Behrens, & Rushworth, 2011; Boorman & Rushworth, 2009; Nicolle et al., 2012; Park, Goïame, O’Connor, & Dreher, 2017), one hypothesis predicts that neural activity in vmPFC/mOFC, and the HC-EC system (**Fig. 1D**), would depend on the current behaviorally relevant dimension alone, while other prefrontal areas may simultaneously reflect the currently irrelevant dimension (**Fig. 1B**). On the other hand, if relationships between people are projected into a unitary space defined by their respective values on two independent dimensions, then we would predict a single neural representation in vmPFC/mOFC and the HC-EC system such that behavioral and neural activity reflect the distance over a 2-D Euclidean space between entities, rather than the behaviorally relevant 1-D rank alone (**Fig. 1C**).

## RESULTS

### Participants learned relational maps of two 2-D social hierarchies and used hubs between them to make inferences between novel pairs of individuals

To investigate whether the human brain constructs a cognitive map of multidimensional social hierarchies and leverages this map for novel inference during decisions, we asked participants to learn the status of unfamiliar people in two separate groups organized hierarchically on two orthogonal dimensions: competence and popularity (**Fig. 1E**). Importantly, participants never saw the 1- or 2-D hierarchies. Instead, they were able to learn the relative ranks of neighboring people who differed by only one level on one dimension at a time through a series of feedback-based dyadic comparisons (see (Kumaran, Melo, & Duzel, 2012)). Participants could then infer the relative ranks of non-neighbors in their group through transitive inference. To learn the two hierarchies, participants completed three days of behavioral training, with a 48-hour gap between each training session, and on the last day, the fMRI experiment (**Fig. 1G**).

During the first two days of training, participants learned the relative status of two groups of 8 “entrepreneurs” separately, on only one dimension per day (**Fig. S1A** for Day 1 and **Fig. S1B** for Day 2 training), such that people in a group were only compared against others belonging to the same group (two groups of 8 entrepreneurs, **Fig. 1E**). At the end of Day 2, test trials without feedback ensured subjects could make transitive inferences to determine the status of remaining members within a group, on each dimension separately (test2 in **Fig. S1B**), indicating they had learned the two 1-D hierarchies for both groups. Importantly, participants were never asked to combine the two dimensions in either group. We included four rank levels per dimension to ensure that differences between rank levels 2 and 3 could not simply be explained by differences in win frequency, since these people each “won” and “lost” on ½ of trials. For the third day of training, fMRI participants learned from select between-group comparisons for the first time (**Fig. S2A**). That is, participants only learned the relative rank of selected entrepreneurs in each group referred to as ‘hubs’, who were paired against both group members (**Fig. S2B and S2C**). By limiting between-group comparisons only to hubs, we were able to create comparative paths connecting each of the individuals in different groups, which could be leveraged to perform model-based inferences between novel pairs of entrepreneurs between groups.

We analyzed fMRI data acquired from twenty-seven subjects who successfully learned the relative ranks of entrepreneurs in each of the two social hierarchies (> 85% performance criterion for inferring relative status of each group member in both dimensions, tested on Day 2 training). To ensure that this behavioral training procedure was sufficient to construct a map-like representation, a separate behavioral experiment consisting of a placement task and a ratings task after Day 3 training, conducted on a separate group of participants, showed that they had successfully learned the four levels for each dimension in the social hierarchy, and importantly, could accurately differentiate between rank levels 2 and 3 for both dimensions (**Fig. S5**). To establish this was also true for our fMRI subject sample, we confirmed that fMRI participants were also able to choose the superior rank face between rank levels 2 and 3 for within-group comparisons: 92.87±0.89% accuracy (t_26_=49.13, p<0.001, one-sample t-test). This performance was not different from other pairs (level 1 *vs*. 2 and level 3 *vs*. 4) also having one-level rank difference (F_2,78_=0.66, p=0.52, one-way ANOVA).

In each trial of an fMRI block, participants were asked to make a binary decision about who was higher rank in one *or* the other dimension between the first face (F1) and the second face (F2) presented sequentially (**Fig. 1F**). Unbeknownst to participants, individuals who were presented at the time of F1 and F2 were selected from different groups and were not hubs in the given dimension (non-hubs; **Fig. S2D**), meaning they had not been previously compared. Following decisions, a third face (F3) was presented, and participants were asked to perform a cover task to simply indicate their gender. F3 was presented from among hubs (**Fig. S2E**) to test for hypothesized fMRI suppression of the relevant latent hub, relative to other matched but non-relevant hubs, that may have been retrieved from memory to guide model-based inferences.

Participants never learned the relative rank of F1 and F2. Instead, they could make a model-based inference through one of two specific hub individuals who had previously been paired with both F1 and F2 during training. Successful inferences, therefore, could rely on building an internal representation of the social hierarchies and a relational memory of the relative positions of F1, F2, and the hub. We predicted that the inference is made along a trajectory in abstract space connecting the two individuals via the (unseen) hub. The hubs were two individuals (H1 and H2) who had been paired with both individuals (F1 and F2) in a task-relevant dimension. The H1 (H2) is uniquely paired with F1 (F2) in between-group comparisons and belongs to the same group with F2 (F1). These task-relevant hubs in one dimension differ from those in the other dimension, which means that to make an accurate inference of the relative status of the same pair of individuals in the two different dimensions, participants needed to retrieve different hubs, which would alter the inference trajectories (e.g. **Fig. S2F**). Since the inference trajectory is anchored by the position of the hub, we were able to track the putative trajectory used by participants by examining which hub between H1 and H2 was selectively retrieved during inferences. Furthermore, we could examine whether participants utilize only the task-relevant rank distance (D) or also the Euclidean distance (E) between individuals’ positions in the cognitive map.

Participants were able to successfully infer the relative position of novel pairs of individuals (overall accuracy ± standard error mean (s.e.m.) = 93.6±0.77%). Nonetheless, the shorter the distance between individuals, the more difficult the decision about relative positions in the hierarchy. To examine the effects of distance of potential trajectories on decision making, we regressed choice reaction times (RT) on different distance measures using a multiple linear regression model, thereby allowing them to compete to explain RT variance. We included the Euclidean distance from the hub (H2) to F1 (E_H2F1_), the Euclidean distance from the other hub (H1) to F2 (E_H1F2_), the relative rank in the task-relevant dimension, which is the 1-D distance between H2 and F1 (D_H2F1_), the 1-D distance between H1 and F2 (D_H1F2_), (**Fig. 1H**), as well as both 1-D and 2-D distances between F1 and F2 (D_F1F2_ and E_F1F2_, respectively), (**Fig. 1I**) to control for their possible covariation with hub-related distances. We found that the greater the 1- D and 2-D Euclidean distance between F1 and H2 (D_H2F1_ and E_H2F1_, respectively), the faster the RT (*β*D_H2F1_±sem=−52.9±11.9, t_26_=−4.5, p=4.5e-05; *β*E_H2F1_±sem=−49.4±5.7, t_26_=−8.8; p=0.003), in addition to an effect of the 1-D distance between F1 and F2 (D_F1F2_) (*β*D_F1F2_±sem=−64.6±12.5, t_26_=−5.3, p=0.0002), (**Fig. 1J** and **Fig. S3A**). Moreover, we found that E_H2F1_ accounted for variation in RTs better than E_H1F2_ (t_26_=−2.73, p=0.01, paired t-test), which did not show a significant effect on RT (*β*E_H1F2_=13.5±5.5, t_26_=1.0, p=0.33). We found that the effects of E_H2F1_ were not different for the trials in which either or both of F1 and F2 was at the highest or lowest rank (i.e. boundary ranks) in the hierarchy compared to the other trials (t_26_=−0.53, p=0.60, paired t-test), and there was a significant effect of the distance for both non-boundary (t_26_=−7.68, p= 3.7e-08) and boundary trials (t_26_=−6.36, p=9.8e-07). We performed an additional multiple linear regression in which the 1-D distances in task-irrelevant dimension (I) were entered as an alternative regressor instead of E (due to the collinearity between E and the sum of D and I). Because E is factorized with two orthogonal vectors, D and I, this analysis allowed us to examine the effects of the D and I without potential collinearity issues between regressors. We found the effects of both 1-D distances from H2 (D_H2F1_ and I_H2F1_), consistent with the finding that E_H2F1_ explains variance in RT over and above D_H2F1_ (**Fig. S3E**). Our behavioral results show that participants preferentially recall H2 as the task-relevant hub to aid in the comparison between novel pairs of faces, with the Euclidean distance to H2 explaining variance over and above the 1-D distance alone.

### Neural activity reflects the Euclidean distance to the retrieved hub during inferences

To examine whether neural activity during choices was likewise modulated by the distance of inference trajectories via the hub over the Euclidean space, first, we regressed BOLD activity at the time of the inference (F2) against the parametric regressors of Euclidean distance between the hub and the target face (E_H2F1_ and E_H1F2_) and their (cosine) vector angles (A_H2F1_ and A_H1F2_). We first tested for effects in *a priori* regions of interest (ROI) that combined multiple anatomically defined ROIs, including the bilateral HC (Yushkevich et al., 2015), EC (Amunts et al., 2005; Zilles & Amunts, 2010), and vmPFC/mOFC (F. X. Neubert, Mars, Thomas, Sallet, & Rushworth, 2014) (**Fig. 1D**). We found neural correlates of Euclidean distance of the decision trajectory via hub H2 (E_H2F1_) in the vmPFC/mOFC (peak voxel [x,y,z]=[2,44,-10], t_26_=4.71 for right vmPFC/mOFC; [x,y,z]=[-2,28,-4], t_26_=4.28 for left vmPFC/mOFC), and bilateral EC (at the peak level, [x,y,z]=[24,-20,-26], t_26_=3.81 for right EC; [x,y,z]=[-18,-10,-26], t_26_=3.49 for left EC) corrected for multiple comparisons over the combined anatomical ROI using permutation-based Threshold-Free Cluster Enhancement (TFCE) (Smith & Nichols, 2009) (p_TFCE_<0.05). We did not find significant effects in HC (p>0.005, uncorrected).

To examine coding outside of our *a priori* ROIs, we performed an exploratory whole-brain analysis. These analyses show that in addition to the EC and vmPFC/mOFC, the right lateral OFC (lOFC, [x,y,z]=[30,34,-18], t_26_=4.12,) activity reflects the Euclidean distance to the context-relevant latent hub H2 to guide inference decisions (**Fig. 2A**). Note that for exploratory analyses, we apply whole-brain TFCE corrections at the threshold p_TFCE_<0.05 (see **Table S1A** for a full list of brain areas surviving TFCE correction at p_TFCE_<0.05). No significant effects were found for the alternative metric terms including the vector angle A_H2F1_, or for metrics associated with the alternative hub, H1 (E_H1F2_ and A_H1F2_) at these thresholds (**Fig. 2A** and **Fig. S7**). These analyses show that activity in vmPFC/mOFC and the EC, but not HC, reflects the Euclidean distance of the trajectory via Hub 2 (E_H2F1_), but not Hub 1 (E_H1F2_), consistent with choice behavior.

**Figure 2.**
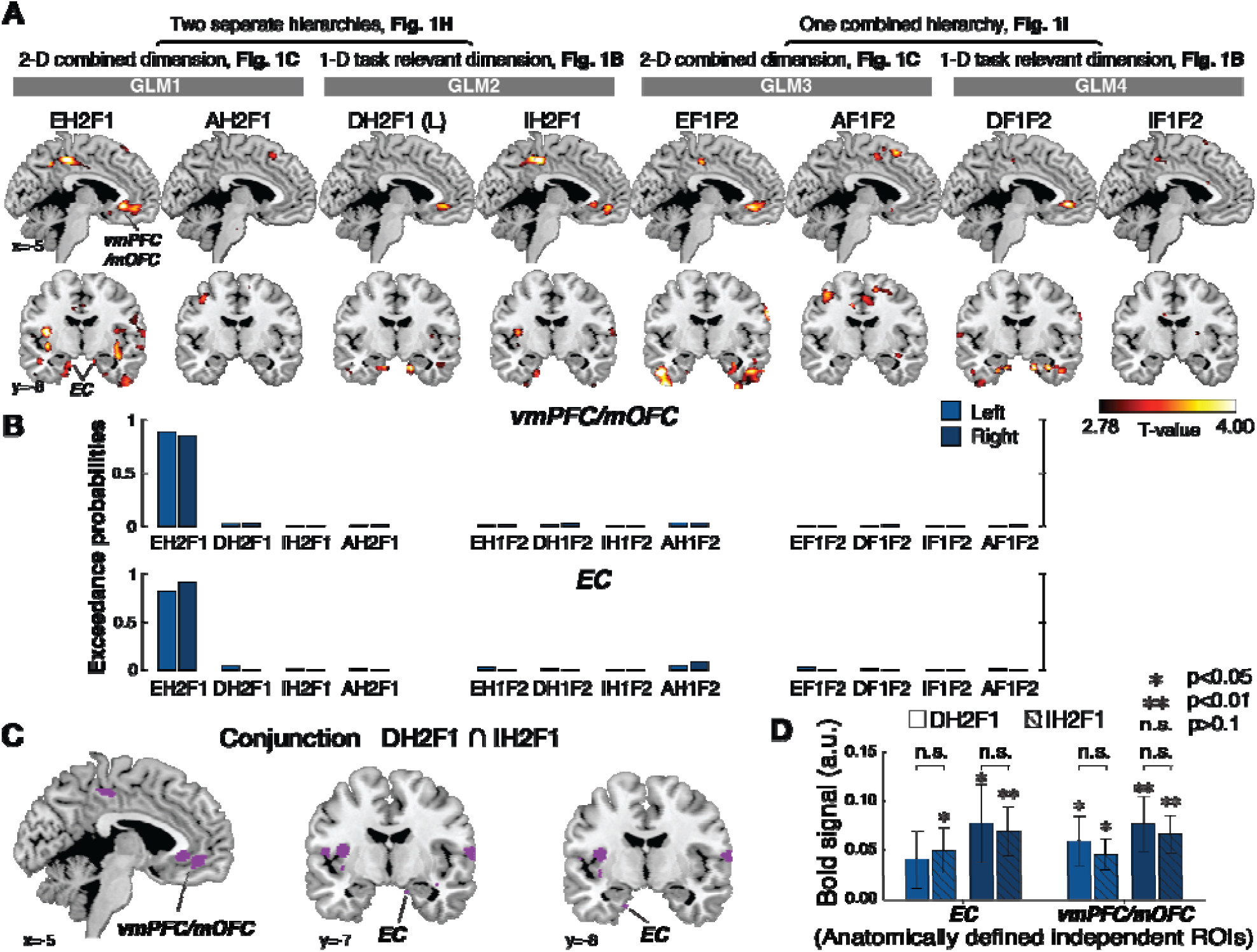
**A**. The bilateral entorhinal cortex (EC) and ventromedial prefrontal cortex (vmPFC/mOFC), p_TFCE_<0.05 corrected within a small volume ROI encode the Euclidean distance from the hub inferred from F2 (H2) to F1 in the 2-D social space (E_H2F1_). Whole-brain parametric analyses showing neural correlates of each of the distance metrics that could theoretically drive inferences between pairs at the time of decisions (F2 presentation). D: 1-D rank distance in the task-relevant dimension (D_H2F1_ and D_F1F2_); L: the shortest link distance between F1 and F2 (L equals to D_H2F1_+1); I: the 1-D rank distance in the task-irrelevant dimension (I_H2F1_ and I_F1F2_); A: the cosine vector angle (A_H2F1_ and A_F1F2_). For visualization purposes, the whole-brain maps are thresholded at p<0.005 uncorrected. **B**. The results of Bayesian model selection (BMS). The exceedance probabilities revealed that the Euclidean distance from the hub (E_H2F1_) best accounted for variance in both EC and vmPFC/mOFC activity compared to the other distance measures, providing evidence that these regions compute or use a Euclidean distance metric to a retrieved hub (H2) in abstract space in order to infer the relationship between F1 and F2. **C**. The conjunction analysis using the minimum statistic at t_26_>2.78, p<0.005 (Nichols et al., 2005) shown in purple revealed that both D_H2F1_ and I_H2F1_ are both reflected in the vmPFC/mOFC and the EC bilaterally. **D**. The activity encoding D_H2F1_ does not differ from that encoding I_H2F1_ in the EC and vmPFC/mOFC, even at a lenient threshold (p>0.1), suggesting that these areas encoding E_H2F1_ assign equal or similar weights to D_H2F1_ and I_H2F1_ (consistent with activity reflecting E_H2F1_) during decision-making.

Next, we investigated our competing hypothesis that the brain flexibly switches between behaviorally relevant and irrelevant dimensions with simultaneous coding of both dimensions, but in different brain regions. Specifically, we tested whether the current behaviorally relevant rank distance (D_H2F1_) and the behaviorally irrelevant rank distance (I_H2F1_) better explain neural activity in the same ROIs, or elsewhere in the brain (GLM2). This analysis revealed positive effects of both D_H2F1_ and I_H2F1_ in vmPFC/mOFC (**Fig. 2A** and **Table S1B**) (D_H2F1_: [x,y,z]=[-2,32,-6], t_26_=5.25, and [x,y,z]=[4,30,-6], t_26_=4.96, and I_H2F1_: [x,y,z]=[6,36,-6], t_26_=3.73, [x,y,z]=[-4,30,-4], t_26_=3.48) (p_TFCE_<0.05). We found similar effects in the EC (D_H2F1_: [x,y,z]=[24,-20,-32], t_26_=3.84 and I_H2F1_: [x,y,z]=[22,-14,-42], t_26_=3.46) at the uncorrected threshold of p<0.001, but they did not survive TFCE correction. Consistent with our analysis of E_H2F1_, we did not find any effects of D_H2F1_ and I_H2F1_ in the HC even at a reduced threshold (p>0.005, uncorrected). To test whether any areas preferentially encoded task-relevant (D_H2F1_) or irrelevant (I_H2F1_) distances, we also directly contrasted these distance terms. Effects in the vmPFC/mOFC and EC were not significant when contrasting D_H2F1_ over I_H2F1_ and I_H2F1_ over D_H2F1_ (**Fig. S4A**). Importantly, we did not find evidence to support the hypothesis that the task-relevant distance (D_H2F1_) was encoded in one set of brain regions and the task-irrelevant distance (I_H2F1_) was simultaneously encoded in a different set of brain regions, even at a liberal threshold (p<0.01, uncorrected) (**Fig. S4A**).

To examine whether the brain preferentially encodes the Euclidean distance of the decision trajectory (E_H2F1_) over and above the rank difference in the 1-D social hierarchy (D_H2F1_), we conducted several additional analyses (see methods for details). First, we confirmed the effect of E_H2F1_ in vmPFC/mOFC ([x,y,z]=[6,42,-14], t_26_=3.75, and [x,y,z]=[-12,24,-20], t_26_=3.72) and EC ([x,y,z]=[30,-14,-30], t_26_=3.35), even after partialling out the 1-D task-relevant distance, D_H2F1_ (p_TFCE_<0.05) (**Fig. S4B**). Second, if the vmPFC/mOFC and EC reflect E_H2F1_, we would expect to find the effects of D_H2F1_ and I_H2F1_ in the same voxels (though the effects of D_H2F1_ and I_H2F1_ would be expected to be weaker compared to E_H2F1_ because E_H2F1_ is factorized into vectors D_H2F1_ and I_H2F1_ and each of these only partially explain the variance in E_H2F1_). Note, the objective of this analysis is to examine what combination of D and I are reflected in vmPFC/mOFC and EC activity, rather than test the hypothesis that these areas independently code for both D and I. If a brain area encoding E_H2F1_ assigns equal or similar weights to D_H2F1_ and I_H2F1_ during decision-making, we would expect that a conjunction null analysis (Nichols, Brett, Andersson, Wager, & Poline, 2005) would reveal overlapping effects of D_H2F1_ and I_H2F1_. Using the minimum statistic conjunction approach, we found inclusive masking between D_H2F1_ and I_H2F1_ (at t_26_>2.78, p<0.005) in the vmPFC/mOFC and EC (**Fig. 2C**). Collectively, the results of these analyses support the interpretation that vmPFC/mOFC and EC activity encodes or reflects E_H2F1_, which is composed of similar weighting of D_H2F1_ and I_H2F1_ (**Fig. 2D**), during novel inferences, consistent with a direct inference over the 2-D space (see **Fig. 1A and 1C**).

Finally, to formally arbitrate between different possible decision trajectories, we used Bayesian model selection (BMS) to compare 2-D and 1-D metrics for different possible trajectories (or comparisons) through the hub H1 (E_H1F2_, D_H1F2_, and I_H1F2_), those through the hub H2 (E_H2F1_, D_H2F1_, and I_H2F1_), and also direct distances between F1 and F2, rather than trajectories via the hub (E_F1F2_, D_F1F2_, and I_F1F2_). In addition, different hypothetical cognitive spaces may have different underlying metrics. That is, if participants do not construct a cognitive map in a 2-D Euclidean space, but rather a different architecture for representing social hierarchies, alternative distance metrics may better account for the neural data than the Euclidean distance. For example, if inferences were made through the sequential retrieval of individuals linking F1 to F2, activity in EC and vmPFC/mOFC should be better explained by the shortest number of links (L; note this is equivalent to 1+D_H2F1_). Alternatively, if a cognitive map encodes only the vector angle between individuals in a polar coordinate system, neural activity should encode the angle between individuals (A_F1F2_, A_F1H1_ and A_F2H1_), while it should be invariant to the length (**Fig. S6**). This formal comparison revealed clear evidence in favor of the Euclidean distance through hub H2 (E_H2F1_) in the EC and vmPFC/mOFC, supporting the hypothesis that the relevant latent hub H2 is used for model-based inference using a cognitive map in Euclidean space (exceedance probability=0.82 in left EC; 0.91 in right EC; 0.89 in left vmPFC/mOFC;0.85 in right vmPFC/mOFC; **Fig. 2B**; **Table S2**). Taken together, our findings show that EC and vmPFC/mOFC compute or utilize Euclidean distances over the 2-D social space to guide inference decisions.

### HC reinstates the hub to guide inferences

The behavioral and neural analyses presented so far provide independent and convergent evidence that the context-relevant hub is retrieved from memory to guide inferences. We therefore searched for neural evidence of a reinstatement of the latent hub along this trajectory to guide decisions. Given the well-established role of the HC in episodic memory retrieval (Diana, Yonelinas, & Ranganath, 2007; O’Reilly, Bhattacharyya, Howard, & Ketz, 2014), we predicted that the HC specifically would reinstate the context-relevant hub to guide inferences between two faces that had never been compared before. To address this question, we adopted a variant of repetition suppression (RS), but for a retrieved rather than explicitly presented item. Notably, RS has been proposed as a means to assess the information content of neuronal ensembles in the human brain (Barron, Garvert, & Behrens, 2016; Boorman, Rajendran, O’Reilly, & Behrens, 2016; Grill-Spector, Henson, & Martin, 2006; Klein-Flugge, Barron, Brodersen, Dolan, & Behrens, 2013). During F3 presentation, participants were exposed to one of eight hub individuals (**Fig. S2E**). We hypothesized that if the relevant hub that bridges F1 and F2 in the given dimension is presented during F3 presentation, directly after participants retrieve the relevant hub, then the BOLD signal in areas reinstating that hub should be suppressed compared to other trials presenting matched but non-relevant hubs. We included only hubs as F3 because these individuals are equally matched for win/loss frequency (each winning on ½ of trials and losing on the other ½) and experience (i.e. presentation frequency), thereby ruling out these potential confounding factors. Moreover, we ensured the Euclidean distance from F2 to F3 (E_F2F3_) was not different when F3 was H1, H2, or a non-relevant hub (F_2_=0.77, p=0.47, one-way ANOVA; **Fig. S3C**), in order to control for the distance between presented faces for each type of hub.

We found that the right HC (peak voxel [x,y,z]=[38,-22,-12], t_26_=3.41, p_TFCE_<0.05 corrected in an anatomically defined bilateral HC ROI (Yushkevich et al., 2015)) showed greater suppression, specifically for the relevant H2 presentations (*β* =−0.46±0.13) compared to all non-relevant hub presentations (*β*=−0.19±0.11) (t_26_=4.54, p<0.001, paired *t*-test in the independent, anatomically defined ROI). The activity in the right HC differed significantly according to which type of hub was shown at F3 presentation (Wilks’ *λ*=.553, F_2,25_=10.11, p=0.001, repeated-measures ANOVA in the independent, anatomically defined ROI). *Post-hoc* paired t-tests between conditions showed a significant effect specific for the relevant H2 compared to all non-relevant hubs (*β*=0.27±0.06; t_26_=4.54, p<0.001), but not between H1 and non-relevant hubs (*β*=−0.18±0.22, t_26_=−0.81, p=0.43). Differences between H2 and H1 were marginally significant (*β*=−0.44±0.23, t_26_=1.95, p=0.06). There was no significant difference in the level of suppression between the right and left HC effects (mean difference *β*=−0.06±0.03, t_26_=−0.45, p=0.66, paired *t*-test), (**Fig. 3**). Furthermore, in an exploratory whole brain analysis, we confirmed that the right HC is the only brain area showing this suppression effect at this threshold. We did not find any brain area showing greater suppression during presentation of the other possible hub, H1 (*β* =− 0.02±0.21 in the right HC; t_26_=0.81, p=0.43), consistent with our analyses reported above, indicating that participants wait for the presentation of F2 to make a backward inference about its rank relative to F1 by preferentially retrieving H2.

**Figure 3.**
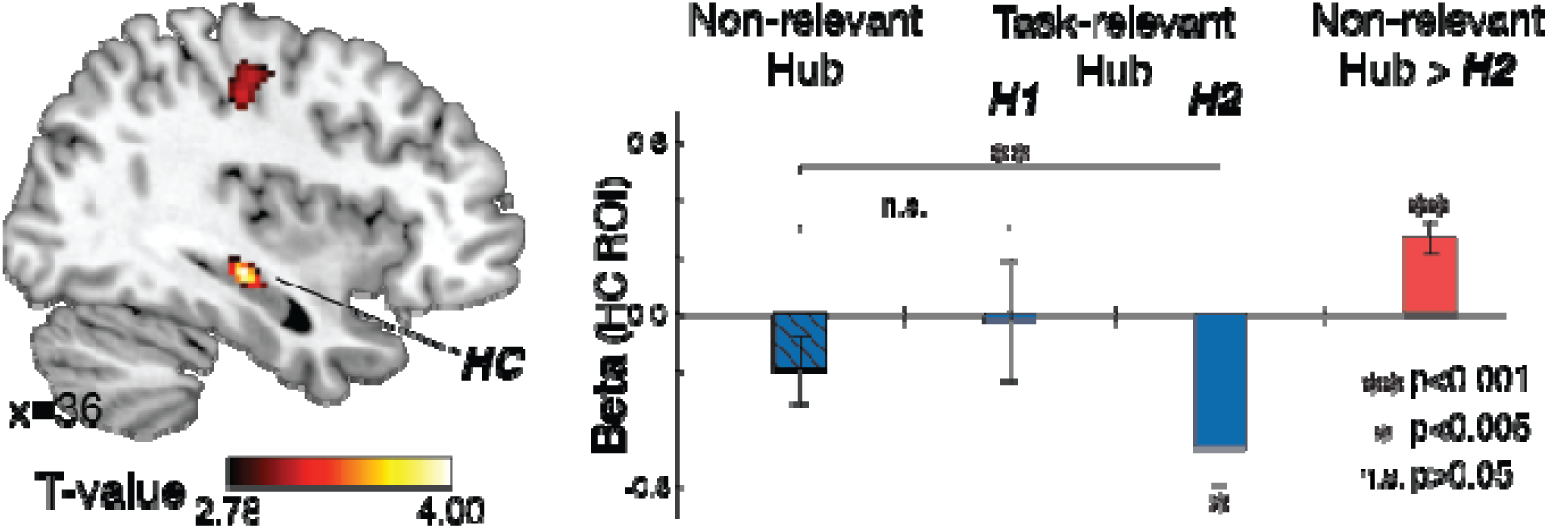
The result of repetition suppression analyses when one of the eight hubs was presented randomly following F2 presentation, as subjects performed a cover task (F3 presentation). Left: BOLD contrast of task-irrelevant hub (Non-relevant hub) > H2, displayed at p<0.005 uncorrected (no masking is applied to the image). The hippocampal (HC) effect is significant at p_TFCE_<0.05 corrected in an independent anatomically defined bilateral HC ROI (Yushkevich et al., 2015), t_26_=3.81, [36,-24,-10]). Right: beta estimates from an independently defined right HC ROI (see **Fig. 1D**). The activity in the right HC differed significantly according to which type of hub was shown at F3 presentation (Wilks’ =.553, F_2,25_=10.11, p=0.001, repeated-measures ANOVA). Activity in the right HC was suppressed when the relevant hub (H2) was presented, compared to matched Non-relevant hubs (p<0.001). No suppression was found when the hub inferred from F1 (H1) was presented (p>0.05).

While this result demonstrated that participants reinstated a specific representation of H2 in the HC to make inferences, we further test whether H2 was preferentially reinstated over H1. We estimated the RS effect of H2 as the difference between the activity in the right HC for the trials in which H2 was presented at F3 compared to other trials in which either a non-relevant hub or H1 was presented at F3 and the same for H1. We found that the suppression effect in the right HC was specific to when H2 was presented compared to all the other trials (*β* =0.28±0.06, t_26_=4.58, p=1.01e-04) but not when H1 was presented compared to all the other trials (*β*=−0.23±0.22, t_26_=−1.05, p=0.30). The difference between the H1 and H2 suppression effects was also significant (Δ*β* =−0.51±0.25, t_26_=−2.10, p<0.05). Importantly, we did not find evidence for decreasing (or increasing) activity in the HC when we modeled whole brain activity as a function of Euclidean distance from the hub to F3 (E_H2F1_ and E_H1F2_) at the time of F3 presentation (**Fig. S3D**), further suggesting that HC suppression was specific to the latent hub itself, rather than driven by proximity in the Euclidean space, thus ruling out a distance-based suppression account between presented faces (Garvert, Dolan, & Behrens, 2017). Taken together, these findings show that HC reinstates the behaviorally relevant hub H2 to guide model-based inferences between distinct relational structures.

### HC, EC, and vmPFC/mOFC represent social hierarchies in a 2-D space

To directly examine the cognitive map’s representational architecture, we measured the pattern similarity between different face presentations during F1 and F2. Under the hypothesis that more proximal positions in the cognitive map will be represented by increasingly similar patterns of neuronal activity, we used representational similarity analysis (RSA) to test the extent to which patterns of activity across voxels in the HC, EC and vmPFC/mOFC (**Fig. 1D**) are linearly related to the Euclidean distance between faces in the true 4-by-4 social network. We reasoned that if the cognitive map of the social network is characterized by two independent dimensions projected into a Euclidean space, the level of dissimilarity between neural representations evoked by each face (**Fig. 4A**) should be explained by pairwise Euclidean distances (E) (**Fig. 1C** and **Fig. 4B**), in addition to the pairwise rank differences in the task-relevant dimension (D) (**Fig. 1B** and **Fig. 4B**).

**Figure 4.**
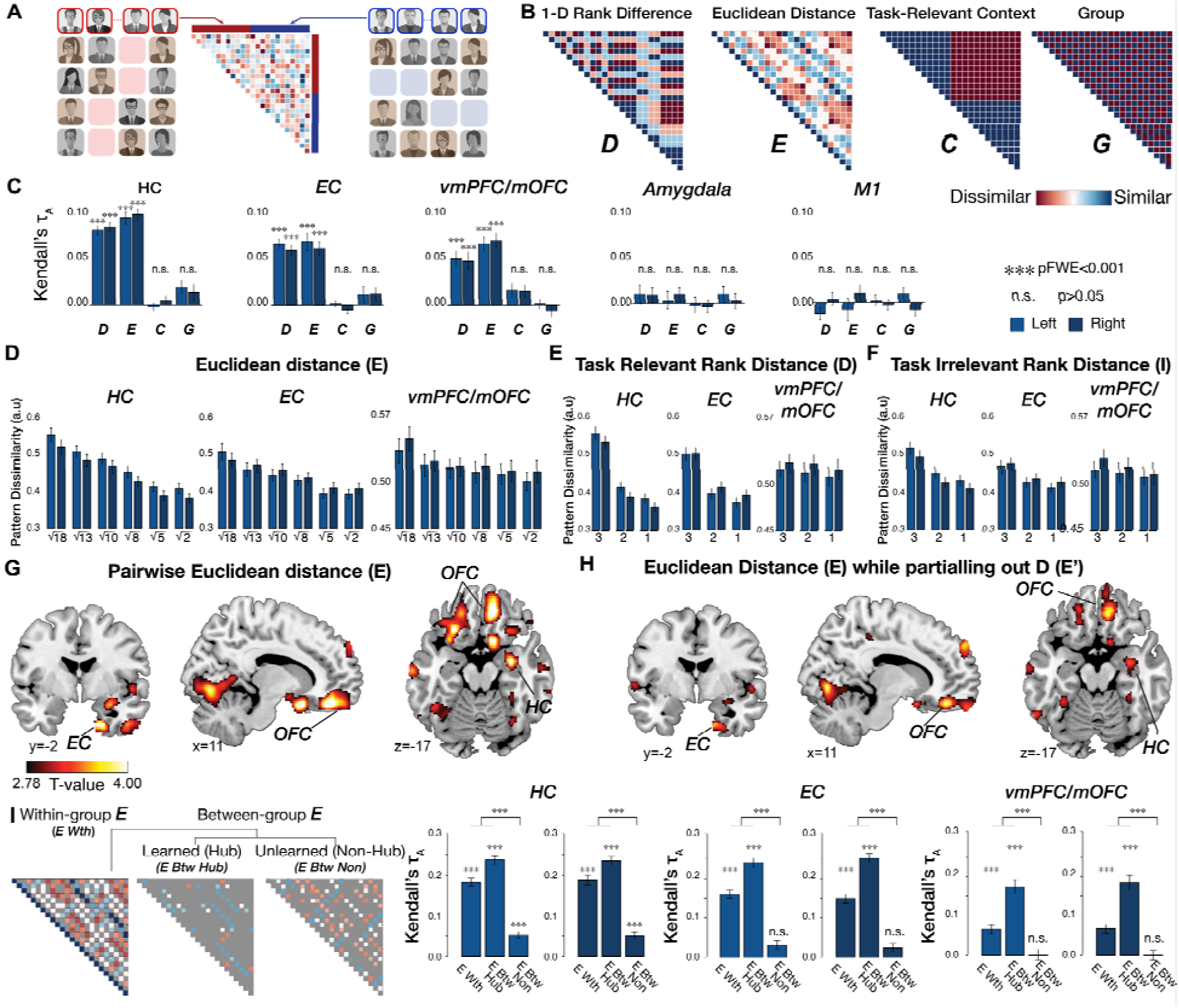
Representational similarity analysis (RSA). **A**. The representational dissimilarity matrix (RDM) was computed in *a priori* regions of interest (ROIs) from the pairwise Mahalanobis distance in the multi-voxel activity patterns evoked when face stimuli were presented at the time of F1 and F2. People were modeled separately when they were shown in the competence (left panel) and popularity contexts (right panel). **B**. The neural RDM was tested against model predictions of four separate dissimilarity matrices, including pairwise differences in the rank in the task-relevant dimension (D), pairwise Euclidean distances on the 2-D social space (E), the behavioral context indicating for which social hierarchy dimension the face was presented (C), and in which group (group 1 or 2) the face belonged during training (G). **C**. Kendall’s τ indicates to what extent a predictor RDM explains the pattern dissimilarity between voxels in each of the ROIs. The model RDMs of D and E, but not C or G, show robust effects on the pattern dissimilarity estimated in the HC, EC, and vmPFC/mOFC (***, p_FWE_<0.001 corrected for the number of ROIs as well as the number of comparisons with the Bonferroni-Holm method). We did not find any significant effects in the amygdala or a control region in the primary motor cortex (M1). **D**. The dissimilarity between brain activity patterns estimated in bilateral HC, EC, and vmPFC/mOFC increases in proportion to the true pairwise Euclidean distance between individuals in the 2-D abstract space. **E and F**. The dissimilarity between activity patterns estimated in bilateral HC, EC, and vmPFC/mOFC increases not only with the rank distance in the task-relevant dimension (D) but also the rank distance in the task-irrelevant dimension (I), suggesting that the HC-EC system utilizes 2-D Euclidean space to represent ranks of individuals (E). **G**. Whole-brain searchlight RSA indicates effects of E in the HC, EC, mOFC (a part of vmPFC), central OFC, and lateral OFC, among other regions (p_TFCE_<0.05). **H**. The activity patterns in the HC, EC, and central and medial OFC are still explained by the model RDM for pairwise Euclidean distance (E) after partialling out its correlation with the model RDM for D (p_TFCE_<0.05; see **Fig. S8C**). For visualization purposes, the whole-brain searchlight maps are thresholded at p<0.005 uncorrected. **I**. The effects of pairwise Euclidean distance (E) between faces and the pattern dissimilarity between voxels in the HC, EC, and vmPFC/mOFC were separately analyzed for within-group (*E Wth*) and between-group relationships (group effects, G). Moreover, the interaction effect between E and G were separately analyzed also based on whether the faces had been directly compared during training (*E Btw Hub*) or not (*E Btw Non*). Effects are strongest for those individuals who had been previously compared during training. That is, activity patterns are better explained by the within-group E and the between-group E of hubs than the between-group E of non-hubs (two-sided Wilcoxon signed-rank test). The between-group E for novel pairs is only significant in HC. Multiple comparisons are corrected with the Holm-Bonferroni method (***, p_FWE_<0.001).

In hypothesis-driven analyses, we first analyzed data from *a priori* selected anatomical ROIs (**Fig. 1D**), including the bilateral HC (Yushkevich et al., 2015), EC (Amunts et al., 2005; Zilles & Amunts, 2010), and vmPFC/mOFC (F.-X. Neubert, Mars, Sallet, & Rushworth, 2015). The representational dissimilarities estimated in the ROIs were explained both by the model representational dissimilarity matrix (RDM) of pairwise Euclidean distance in 2-D space (E, one-sided Wilcoxon signed rank test, df=26, p_FWE_<0.05 Holm-Bonferroni correction for multiple comparisons across numbers of model RDMs (n=4) and bilateral ROIs (n=6)) and, in a separate RDM, by the pairwise rank difference in the task-relevant distance (D, p_FWE_<0.05) between individuals (**Fig. 4C**). Based on previous demonstrations that univariate amygdala activity (Kumaran et al., 2012) and gray matter density (Bickart, Wright, Dautoff, Dickerson, & Barrett, 2011; M. A. P. Noonan et al., 2014; Sallet et al., 2011), correlate with social dominance status, we also tested anatomically defined amygdala ROIs (Tzourio-Mazoyer et al., 2002). The amygdala pattern similarity was neither explained by E nor by D, even at a reduced threshold (p>0.05, uncorrected) (**Fig. 4C**). As a control region, we also tested the pattern similarity in primary motor cortex (M1) (Glasser et al., 2016), which was not explained by either predictor (p>0.05, uncorrected), (**Fig. 4C)**. On the other hand, the pattern similarity in HC, EC, and vmPFC/mOFC was not explained by the behavioral “context” of the task-relevant dimension (C, defined as popularity or competence trials), nor whether individuals belonged to the same group or not during training (G), (p>0.05, uncorrected), (**Fig.4C**; **Table S3A**). Importantly, the predictor of pairwise Euclidean distance (E) still significantly accounted for the pattern similarity in HC, EC, and vmPFC/mOFC (Rank correlation *τ*_A_=0.045±0.005 for HC; *τ*_A_=0.027±0.007 for EC; *τ*_A_=0.048±0.006 for vmPFC/mOFC; p_FWE_<0.001) after partialing out its shared correlation with rank distance (D) to ensure that D alone was not driving the pattern similarity effects (**Fig. S8C**). To confirm that the pattern similarity truly reflected E, we tested for separate effects of D and I. Decomposing E into the terms D (**Fig. 4E**) and I (**Fig. 4F**) revealed a linear relationship between pattern similarity and both distance components (See **Table S3A** for mean rank correlations *τ*_A_; all p_FWE_<0.05 with Holm-Bonferroni correction). These analyses show that, in addition to D, I contributed significantly to representations in these regions, supporting the interpretation that the social hierarchy was represented in 2-D, even though only one dimension was behaviorally relevant.

A natural question arises from this finding: Why do participants need to retrieve the hub for inferences if they have already integrated the two hierarchies into a single cognitive map? We reasoned that if the two 2-D maps, one for each group, had not yet been fully integrated into a single map, then the effect of E should be weakest for different members who had never been compared during training. We found that the effect of E was strongest for within-group pairs (i.e. individuals who were part of the same group during training; **Fig. 4I** and **Fig**.**S9A**) and for between-group pairs involving hubs (i.e. individuals and their hubs who were compared during between-group learning in day 3 training; **Fig. 4I** and **Fig. S9B**), and weakest for never-compared between-group pairs of non-hubs (**Fig. 4I** and **Fig. S9C**) in the bilateral HC, EC and vmPFC/mOFC. Moreover, this difference was significant in all three areas (p_FWE_<0.001, two-sided Wilcoxon signed-rank test; **Fig. 4I**; **Table S3C**; the mean rank correlations *τ*_A_ are shown in **Fig.4I** and **Table S3B**). Notably, however, the effect of E was still significant, though weaker, for never-compared between-group pairs of non-hubs in HC alone, suggesting that HC integration may lead EC and vmPFC/mOFC, since there was no significant effect for these novel pairs in these latter regions. These findings suggest that the previously experienced pairs may have been fully integrated into a single map in each region, but that the novel unlearned pairs were not as accurately integrated, and only present in the HC alone.

An alternative possibility is that some subjects had formed a fully integrated map, but others had not and so these had to retrieve a hub to enable inferences. To test for this possibility, we examined whether individual differences in the level of integration of two social hierarchies explained the different levels of reinstatement of the task-relevant hub across individuals. We found that the levels of hub reinstatement (the size of repetition suppression effects in the right HC specifically to H2 compared to non-relevant hubs) were in fact not explained by either the strength of neural representation (rank correlation, Kendall’s *τ*_A_) of the unlearned relationship itself (between-group of non-hub faces) (r=0.18, p>0.05) (**Fig. S9D**), the relative strength of the unlearned relationship compared to the learned within-group relationship (r=0.18, p>0.05), nor the relative strength of the unlearned relationship compared to the learned between-group relationships (via hubs) (r=06, p>0.05) (**Fig. S9E**). Taken together, these findings suggest that participants have a neural representation that was in the process of combining the two hierarchies. This pattern of findings may explain why participants needed to reinstate the task-relevant hub to make novel inferences before they had completed forming a neural representation of the fully combined hierarchy.

In addition to these hypothesis-driven analyses, we also performed whole-brain exploratory analyses to test whether the neural representation of the social network extends to a broader set of regions. Specifically, we measured the extent to which each predictor (the model RDM of E and D) explains the pattern similarity measured from searchlight-based pattern analyses across the whole brain. This analysis revealed that the pairwise Euclidean distance (E) significantly explained the representational similarity between faces in HC and EC, as shown by the ROI analyses, and also in medial, central, and lateral OFC, among other areas (p_TFCE_<0.05; **Fig. 4G** and **Table S4A**). A separate RDM based on the pairwise 1-D rank distance (D) significantly explained representational similarity between activity patterns in the lateral OFC, medial prefrontal cortex (mPFC), and posterior cingulate cortex (PCC) (p_TFCE_<0.05; **Fig. S8A**; **Table S4B**). Furthermore, partialing out D from the RDM for E revealed significant effects in these same areas of HC, EC, and central/medial OFC, confirming that these representations were not simply driven by D alone (**Fig. 4H**; **Table S4C**). Our findings suggest that the HC, EC and vmPFC/mOFC do not treat dimensions separately when representing individuals in a social network space. Instead, representations vary along the multidimensional cognitive map even when only one dimension is relevant to current behavioral goals.

## DISCUSSION

The HC formation is thought to contain relational codes of our experiences that integrate spatial and temporal dimensions into a multidimensional representation (Buzsáki, 2013; Howard Eichenbaum, 2017b; Howard Eichenbaum & Cohen, 2014; Konkel & Cohen, 2009). Memories of place and their spatial relationship are key elements to constructing a cognitive map of physical space (Butler, Hardcastle, & Giocomo, 2019; Kropff, Carmichael, Moser, & Moser, 2015; Moser et al., 2008). In humans, the ability to construct an accurate cognitive map of relationships between abstract and discrete information is proposed to be critical for high-level model-based decision making and generalization (Behrens et al., 2018; Bellmund et al., 2018; Vikbladh et al., 2019). We show that the HC and EC, which are famously known for their proposed roles in the ability to navigate physical space (Moser et al., 2008; O’Keefe & Nadel, 1978) and simultaneously their roles in episodic memory (H. Eichenbaum, Yonelinas, & Ranganath, 2007; Ekstrom & Ranganath, 2018), contribute in a more general way to the organization and ‘navigation’ of social knowledge in humans (Cohen, 2015; Rubin, Watson, Duff, & Cohen, 2014). Although participants were never asked to combine the two social dimensions, we found that the brain spontaneously represents individuals’ status in social hierarchies in a map-like manner in 2-D space. Such a cognitive map can be used to compute routes through the 2-D space and corresponding distances (Behrens et al., 2018), which we found were computed or used to guide inferences in EC and interconnected vmPFC/mOFC, a region known to be important for value-based decision making (Boorman, Behrens, Woolrich, & Rushworth, 2009; FitzGerald, Seymour, & Dolan, 2009; Hunt et al., 2012; Lim, O’Doherty, & Rangel, 2011; Nicolle et al., 2012; M P Noonan, Mars, & Rushworth, 2011; Maryann P Noonan, Chau, Rushworth, & Fellows, 2017; Papageorgiou et al., 2017; Rushworth, Noonan, Boorman, Walton, & Behrens, 2011; Strait, Blanchard, & Hayden, 2014). Moreover, our results show that the HC-EC system did not selectively represent the task-relevant information in our task, but the relative positions in the multidimensional space. More broadly, these findings support the HC-EC system’s role in representing a cognitive map of abstract and discrete spaces to guide novel inference decisions that relied on that cognitive map.

We found that during novel inferences, RTs and neural activity in EC, vmPFC/mOFC, and lOFC reflected the Euclidean distance of navigational trajectories on 2-D space via relevant hubs, over and above the behaviorally-relevant 1-D ranks. The faster reaction times and the greater EC and vmPFC/mOFC activity also could reflect the certainty in the decision, if it is assumed that the certainty is based only on the 2-D difference between Hub (H2) and F1, as these would be indistinguishable in our task design. It is also possible that when participants use a 2-D social hierarchy representation as a cognitive map, they could use the Euclidean distances between individuals to compute the decision value, which would result in greater certainty about their decision when a longer Euclidean distance guides the decision. Notably, cells encoding the distance to “goals” have been documented in EC and mPFC of rodents (Boccara, Nardin, Stella, O’Neill, & Csicsvari, 2019; Butler et al., 2019; Guise & Shapiro, 2017) when navigating in physical space, while the rodent OFC has been shown to be necessary for model-based inferences (Jones et al., 2012). A separate body of work in humans and monkeys during value-based choice has consistently found value comparison signals thought to guide goal-based choices in vmPFC/mOFC (Boorman et al., 2009; FitzGerald et al., 2009; Hunt et al., 2012; Lim et al., 2011; Nicolle et al., 2012; Papageorgiou et al., 2017; Rushworth et al., 2011; Strait et al., 2014). Taken together, this suggests the EC, vmPFC/mOFC, and lOFC compute or utilize distance computations for inferred trajectories derived from a cognitive map to guide novel decisions.

Notably, the vmPFC/mOFC and EC both contained a neural representation of the 2-D cognitive map, and, during decision making, their activity reflected a direct vector on the 2-D cognitive map from the retrieved hub that guided inference behavior. We interpret these findings as reflecting a decision process that uses a vector over Euclidean space on the cognitive map. We note that, depending on how certainty is defined, this univariate effect may also be interpreted as reflecting the certainty of the decision. This finding is also consistent with an attractor decision-making network whose speed of accumulation to an attractor state will reflect the decision certainty, as has been proposed previously for vmPFC/mOFC in the context of value-based decision making (Hunt & Hayden, 2017; Hunt et al., 2012). That EC also reflects the same term suggests that, unlike in most value-based decision-making studies, it also contributes to the decision computation when based on a cognitive map. Notably, previous findings using human fMRI have reported that EC activity increases with longer Euclidean distances of planned and taken routes and reflects the planned direction of future routes during spatial navigation tasks (Chadwick et al., 2015; Doeller et al., 2010; Howard et al., 2014). Moreover, the global relational codes provided by grid cells can be used for straightforward computation of Euclidean distance from grid fields (Behrens et al., 2018; Bush et al., 2015), and has been interpreted as a mechanism, in combination with other neural codes in EC and HC, for vector navigation to goals during planning (Banino et al., 2018; Behrens et al., 2018). This view suggests that greater EC activation for greater Euclidean distances may reflect this computation for the inferred vectors that guide inferences for non-spatial decision making, in concert with choice selection processes in vmPFC/mOFC.

These trajectories imply that people retrieve the task-relevant hub (H2) to guide backward inferences during novel comparisons between people from different groups. Consistent with this interpretation, suppression analyses revealed that the HC was the only brain region showing a reinstatement of this same unseen hub (H2) at the time of inference decisions, in order to compare two individuals whose positions were learned in different social groups. This effect was specific to H2 (relative to both non-relevant but matched hubs and H1). Importantly, this H2-specific suppression in the HC could not be explained by the Euclidean distances between F2 and F3 (**Fig. S3C**) or H2 and F3 (**Fig. S3D**), nor by win/loss frequency, nor the degree of experience with each face, since all faces presented as F3 were hubs that were carefully matched for presentation frequency/familiarity and win/loss history. Note that making inferences via H2 does not indicate that participants integrated members in one specific group to the pre-existing structure of the other group. Considering that participants were also presented all F1-F2 pairs in reverse order, our results indicate that a different hub was preferentially recalled during inferences about the same pairs of individuals when they were presented in reverse order. Although we cannot rule out the possibility that H1 was also reinstated transiently at the time of F1 presentation, our behavioral analyses of RT (**Fig. 1**), univariate neural analyses of trajectory distances (**Fig. 2**), and suppression analyses (**Fig. 3**) all provide independent and convergent evidence that H2 was reinstated to guide inference behavior. Given the fact that participants were never informed that F1 and F2 were selected from non-hubs in different groups, it appears to be a more natural decision to select the hub to guide inferences after knowing both F1 and F2.

We found that the pattern similarity between faces in HC, EC, and in vmPFC/OFC was robustly and linearly related to the true Euclidean distance between faces in the 4-by-4 social network, such that closer faces in the abstract space were represented increasingly more similarly. This finding is striking for two reasons: first, the two dimensions never had to be combined to perform the task accurately; and second, the true structure was never shown to participants, but had to be reconstructed piecemeal from the outcomes of binary comparisons between neighbors in each dimension separately learned on separate days. There are several strategies that could, in principle, be used to solve this task that do not rely on a 2-D representational space. For example, the task-relevant and irrelevant dimensions could be represented in separate brain areas, a hypothesis for which we did not find support. Alternatively, each person’s rank could have been represented by a linear (or logarithmic) number line, such as that found in the bilateral intraparietal area (Piazza, Izard, Pinel, Le Bihan, & Dehaene, 2004), or as a scalar value that is updated using model-free mechanisms. That the neural representation in the HC, EC, and vmPFC/OFC areas automatically constructed the 2-D relational structure in our tasks instead suggests that the brain may project people, or perhaps any entities, into a multi-dimensional cognitive or relational space such that the entity’s position is defined by the feature values on each dimension (Bellmund et al., 2018; Buzsáki & Tingley, 2018; Howard Eichenbaum, 2017b). It is unclear from our study whether this finding is specialized to representing people, who are likely to be ecologically perceived as coherent entities over time, and characterized by multiple attributes, or more general to representing any entity. Precisely how this construction takes place and its generality will be an important topic for future studies to investigate.

Following recent theoretical proposals (Howard Eichenbaum & Cohen, 2014; J. C. Whittington et al., 2019) inspired by seminal studies (Dusek & Eichenbaum, 1997; Tolman, 1948), we hypothesize that subjects abstract structural representations from the ordinal comparisons about rank in the social hierarchy, which, by virtue of the inferred structure, allows efficient direct inferences to be made from limited observations. One commonly experienced topology in the real world is Euclidian space, so one intriguing possibility is that the brain uses a neural system that represents relationships in continuous spatial problems to represent distances and directions of non-spatial, ordinal (and possibly even discrete) relationships, such as social networks, and, recent evidence suggests, conceptual (Bellmund et al., 2018; Constantinescu et al., 2016; Quiroga, 2012) and even task spaces (Baram, Muller, Nili, Garvert, & Behrens, 2019). We note that it would be difficult to account for the effects of Euclidian distance observed in univariate and multivariate analyses from chaining together previously experienced associations alone. Explicitly representing the hierarchical structure would allow for generalization of learned relationships to other problems that share structure, and also efficient and direct inferences, as we observed (J. C. Whittington et al., 2019). In our task, we examined the form of inferred trajectories by formally comparing the Euclidian distance with the link distance (how many learned associations are travelled through for inferences, which we call L). For univariate effects during decision making, we found evidence supporting E over L in vmPFC/mOFC and EC (see **Fig. 2**). However, it is also possible that non-ordinal relationships would not be represented in a Euclidian space, but instead more like a discrete graph.

Recent studies suggest that even relationships between social entities may be organized as a cognitive map in HC. In particular, these studies suggest that the HC not only represents the position of others in physical space through the firing fields of “social place cells” (Danjo, Toyoizumi, & Fujisawa, 2018; Omer, Maimon, Las, & Ulanovsky, 2018), but also their 1-D rank in a dominance hierarchy, with greater HC BOLD activity for higher rank people (Kumaran, Banino, Blundell, Hassabis, & Dayan, 2016; Kumaran et al., 2012), self-other differences in preference-based ratings (Kaplan & Friston, 2019), and their egocentric relative direction in a space characterized by power and affiliation dimensions (Tavares et al., 2015). This latter study showed that univariate HC BOLD activity is modulated by the angle of the vector to the relative position of another person with whom subjects interacted during a role-playing game (Tavares et al., 2015). Notably, the cosine angle, unlike the Euclidean distance, is invariant to the distance to others. While this important study first suggested the HC encodes the egocentric direction of another’s relative position during online social interactions, the nature of HC’s representational architecture, and any putative role of the EC and vmPFC/mOFC in representing a cognitive map of social space and, critically, how the cognitive map is used to perform novel inferences were previously unaddressed. By showing that the neural activity patterns in the EC and vmPFC/mOFC represent the 2-D social hierarchy and their activity reflects the Euclidean distance of inferred decision trajectories as the decision-value that guides choices, we provide evidence that these networks were engaged in map-guided decision-making. In addition to dominance, theories in social cognition propose orthogonal psychological dimensions of competence and warmth along which humans represent other people in an abstract space (Fiske, 1992). We found that 2-D social hierarchies defined by independent social dimensions were represented and navigated as a cognitive map to guide novel inferences. Our study further elucidates how even social relationships are represented as a cognitive map and how that map guides novel inferences about social status. In so doing, it adds to growing evidence to suggest that the same neural system used for representing non-social relationships may be leveraged to represent and make inferences about social relationships.

Another recent study (Tang et al., 2019) in which human participants learned the association between novel visual stimuli and reward values over multiple days used a linear support vector machine to show that the brains of fast learners were more likely to use an efficient coding scheme to represent stimuli. Their study suggested that multidimensional coding in the brain helps participants to discriminate different stimuli while simultaneously embedding the stimuli in an efficient low-dimensional task-relevant structure. In the current task in which the rank positions of individuals in both dimensions were used to make inferences about the social hierarchical status of novel pairs of individuals, our results suggest that the accurate representation of multidimensional task-relevant information may be critical for successful model-based decision-making.

While we found strong effects of the Euclidean distance between patterns of activity evoked by pairs of people across the entire social network, post-hoc comparisons indicated that these effects were strongest for pairs of faces within the same training group, and for pairs of faces and their previously paired hubs, while present in HC alone, but significantly weaker for novel pairs of faces between groups (**Fig. 4I**). The relatively weaker effects of Euclidean distance specifically for the unlearned relationships are consistent with the interpretation that people had formed two 2-D cognitive maps, one per hierarchy or group, and were in the process of combining these into one map of the integrated social hierarchy. This pattern also dovetails with our finding that people utilized hubs to guide inferences between pairs in different groups, rather than relying solely on the vector distances between faces from different groups. Notably, our findings suggest subjects used both model-based inferences through a retrieved hub, and direct Euclidian inferences from that hub to the within-group face, to make decisions. This interpretation is also supported by a recent computational model suggesting that while place cells in HC supports topological navigation between directly experienced neighbors, grid cells in EC are integral to support metric vector navigation to connect unexperienced pairs (Edvardsen, Bicanski, & Burgess, 2019). Moreover, previous studies have shown that rat grid cells that had already been formed to represent two identical but separate spatial compartments gradually begin to represent the unified combined space when the two compartments are merged by opening a connected corridor (Carpenter, Manson, Jeffery, Burgess, & Barry, 2015) or removing the wall between compartments (Wernle et al., 2018). Both of these studies found incomplete grid fields as the representation reflected a presumed process of integrating the two physical spaces before fully learning the merged space. We speculate that an analogous gradual process of integration is at play in our study, one that would be intriguing to continue to track over time.

As an alternative explanation, there could be individual differences in the retrieval of the hub and the level of integration of the two social hierarchies. While our sample size (n=27) might be too small to test for individual differences robustly, we did not find evidence that individual differences in the level of integration of the two hierarchies explained the tendency to rely more on retrieving the hub to guide inferences (**Fig. S9 D and E**). Notably, this observation of different effects of Euclidean distance in experienced and unexperienced relationships are consistent with the view that the training episode may have constituted a context during which each relational map was initially formed separately for each group in HC, EC, and vmPFC/mOFC (Clewett, DuBrow, & Davachi, 2019; Diana et al., 2007; McKenzie et al., 2014; Sols, DuBrow, Davachi, & Fuentemilla, 2017), supporting the theory that the elements of the map are initially bound together within distinct temporal event contexts.

It is noteworthy that the HC representation was the most flexible, putatively leading EC and vmPFC/mOFC in the integration of the two groups into a single coherent representation. This finding is reminiscent of faster adaptation of place fields than grid fields to new reward locations (Boccara et al., 2019) and changes to an environment’s geometry (Munn, Mallory, Hardcastle, Chetkovich, & Giocomo, 2020). Nevertheless, our results indicated that participants relied on the hub to perform model-based inferences between never compared members of each group using reinstatement of the retrieved hub in HC. It is possible that the integrated HC representation did not appear to be used for direct inferences between groups is because the combined representation was relatively imprecise compared to within-group pairs (**Fig. 4I**), or because it was simply easier for subjects to rely on the recently well learned hub comparisons for inferences. Alternatively, that subjects did not appear to rely on these integrated HC representations may suggest that coherent representations of the combined map must also form in EC and perhaps vmPFC/mOFC, a process that requires more learning or time, to perform vector navigation.

Our findings suggest that the brain utilizes the same neural system for representing and navigating continuous space to code the relationship between discrete entities in an abstract space. Further, they suggest that accurate inferences about relative ranks of novel pairs of individuals may depend on the ability to find a direct route in a multidimensional space. This vector-based navigation over the cognitive map may be critical for efficient decision making and knowledge generalization. Moreover, accurate knowledge about the position of others in a social space should provide a solid foundation for sound inferences, thereby supporting effective model-based decision making. We found that the same cognitive map constructed by the HC-EC system is present in other brain areas, including the interconnected vmPFC/mOFC (Barbas & Blatt, 1995; Howard Eichenbaum, 2017a; Insausti & Muñoz, 2001; Preston & Eichenbaum, 2013; Wikenheiser et al., 2017) and neighboring central/lateral OFC, generally supporting the theory that OFC represents a cognitive map of task space (Schuck et al., 2016; Wikenheiser & Schoenbaum, 2016; Wilson et al., 2014), though in our study not only of the behaviorally relevant task space, but also the broader task space. Moreover, we show here that the OFC’s representation of the task space respects map-like Euclidean distances of vectors through that space and that OFC activity reflects these distances to latent hubs retrieved from memory to guide inference. These findings thus cast light on why the OFC plays a critical role in model-based inference on the one hand (Jones et al., 2012), and damage to the HC-EC system also impairs complex, model-based decision making on the other (Gupta et al., 2009; Miller, Botvinick, & Brody, 2017; Vikbladh et al., 2019).

Finally, we suggest that the HC-EC system may play a key role in constructing a global map from local experiences, which may guide model-based decisions in vmPFC/mOFC, a region previously implicated in value-guided choice (Boorman et al., 2009; Chib, Rangel, Shimojo, & O’Doherty, 2009; Grabenhorst & Rolls, 2011; Hunt et al., 2012; Lim et al., 2011; Papageorgiou et al., 2017; Strait et al., 2014). This same cognitive map appears to further guide how humans integrate knowledge in the social domain, a critical ability for navigating our social worlds.

## METHODS

### Participants

A total of 33 participants (16 female, age range: 19–23, normal or corrected to normal vision) were recruited for this study via the University of California, Davis online recruitment system. Six participants were excluded due to strong head movements larger than the voxel size of 3□mm. In total, 27 participants entered the analysis (mean age: 19.37±0.26, standard error mean (SEM)). The study was approved by the local ethics committee, all relevant ethical regulations were followed, and participants gave written consent before the experiment.

### Stimulus

The stimuli consisted of 16 grayscale photographic images of faces (Strohminger et al., 2016) and two colored cues (red and blue squares). Each of the colored cues indicated the task-relevant dimension of the social hierarchy for the current trial. The red square indicated the competence hierarchy for one-half of participants and the popularity hierarchy for the other half. All images were adjusted to the same mean grayscale value. The inter-trial fixation target was a white cross in the middle of a black screen. For hub learning and the fMRI experiment, the inter-stimulus fixation target was a purple cross (between F1 and F2) and a green cross (between F2 and F3) in the middle of a black screen, which indicates the progress of each trial to participants. The stimuli were presented to participants through a mirror mounted on the head coil. Note that face stimuli presented in this paper are license-free images for display purposes. Prior to the experiment on the first day of training, participants performed a 1-back task where they viewed each individual face three times to minimize stimulus novelty effects.

### Social hierarchies

Participants were asked to learn the relative ranks of 16 individuals (face stimuli) in two dimensional social hierarchies defined by popularity and competence. The 16 face stimuli were introduced as entrepreneurs; participants were asked to learn about which individuals were more capable to attract crowd funds (labelled popularity) and which individuals had higher technical proficiency (labelled competence) and used this information to guide investment decisions.

Each hierarchy has four levels of ranks. Four individuals were allocated at the same rank at each level of the hierarchy. Therefore, the structure of multidimensional social hierarchies is 4×4 (**Fig. 1E**). The rank of an individual in one dimension was not related to his/her rank in the other dimension. For instance, the rank of four individuals who are at the 1^st^ rank in the popularity dimension are the 1^st^, the 2^nd^, the 3^rd^, and the 4^th^, respectively in the competence dimension. During first two days of training, the relative status of one individual is only compared to one-half of the other face stimuli, which implicitly creates two groups in which each group comprised eight individuals (**Fig. S1**). In each group, two individuals were allocated into the same rank of each of the dimensions (2 × 4 ranks = 8 individuals). The sub-group structure is shown in **Fig. 1E** and **H**. The allocation of each face to the position in the social hierarchies was pseudo-randomized, in order to make sure that any visual features of the face (gender, race, and age) were not associated with the rank of the individuals. To do this, we prepared eight stimuli sets. Each of the stimuli sets comprise 16 different faces. A stimuli set was randomly assigned across participants.

### Task instruction and experiment procedures

Participants were instructed to imagine that they were a venture capitalist and decide where to invest after learning the relative ranks of 16 entrepreneurs in two independent dimensions − competence and popularity. Participants were asked to learn which individual was better in technical proficiency – competence hierarchy, and which individual was better in attracting crowd funding – popularity hierarchy. During the learning block, participants were presented with two face stimuli of entrepreneurs with a contextual cue indicating the task-relevant dimension, chose the higher rank individual in the given dimension, and received feedback at the end of every trial. Participants were told that they would need to use the knowledge acquired during the learning block to decide in which entrepreneur they would want to invest when the ability in only one social hierarchy dimension is important. During the test block, participants chose one of two face stimuli who they believed was higher in the given dimension. They did not receive any feedback during test block trials.

Before training, the following instructions were clearly given to participants: (1) two entrepreneurs presented in the learning block have one rank difference whereas two entrepreneurs presented in the test block can have one or more rank differences. (2) Multiple individuals could be allocated to the same rank. Importantly, participants were never given any information implying the structure of social hierarchies, such as the total number of ranks in each dimension, or the number of individuals allocated into the same rank and were never asked to solve the task spatially. Each subject participated in behavioral training across three separate days, at least 48 hours apart. After the behavioral training on the third day, subjects participated in the fMRI experiment.

### Behavioral training

The behavioral training comprised ‘learning’ blocks and the ‘test’ blocks (**Fig. 1C**). Training began with learning block mini-blocks. In the beginning of each of the mini-blocks, participants were presented which block they were in (competence or popularity). The purpose of the learning blocks was to provide an experimental setting in which participants would gradually acquire knowledge of two social hierarchies (one per group) through piecemeal experiences of comparisons for pairs with only one rank-level difference.

For the test blocks during training, participants were asked to infer the relative rank between any two individuals. To perform correctly, therefore, participants need to make successful transitive inferences. If they adopted an alternative learning strategy, such as model-free learning – i.e. comparing the values assigned to each of the face stimuli according to their number of wins/loses − participants should not be able to distinguish the second and third rank individuals, since their number of wins/loses were equal (though see (Frank et al., 2005)). Participants who successfully distinguished the second and third rank individuals above chance while also reaching above 85% accuracy overall in each test block were invited to continue participating in the fMRI experiment.

It is important to note that, during three days of training, each of the 16 individuals was presented the same number of times to participants. For trials in which participants responded too slowly (>2s), feedback was not given and the missing trial was tested again after a random number of trials to ensure all participants could in principle acquire the same level of knowledge. Importantly, participants were never asked to combine individuals’ ranks in both dimensions to make decisions, and they were never shown either the one-dimensional (1-D) or two-dimensional (2-D) social hierarchies.

#### Learning blocks of day 1 and day 2 training: Learning relative ranks of within-group members

During learning blocks, participants were presented two face stimuli having one rank difference on a black screen with a colored contextual cue indicating which was the task-relevant dimension in the current trial (learning block in **Fig. S1A**). They were asked to indicate who was superior in the given social dimension. Participants learned the relative status of all possible one rank difference pairs with feedback in random order. Feedback (correct/ incorrect) followed at the end of all responded trials. For the learning block of day 1 training, participants learned the relative status of an eight-individual group in one of two social hierarchies for the first mini-block (e.g. the hierarchy in the competence dimension for group 1 individuals). For the second mini-block, they learned the relative status of the other eight-individual group in the other social hierarchy (e.g. the hierarchy in the popularity dimension for group 2 individuals). For the learning block of day 2 training (the learning block in **Fig. S1B**), they learned the relative status of each group individuals in the unlearned hierarchy dimension (e.g. the hierarchy in the popularity dimension for group1 individuals and the competence dimension for group2 individuals). After two days of training, in principle participants could have learned the two different 2-D social hierarchies (one per group). The right panel in **Fig. S1** shows the hypothesized structure of the cognitive map that participants could have built at the end of each training day. For both day 1 and day 2 training, participants completed eight mini-blocks in the learning blocks (**Fig 1G**). One-half of participants learned the relative ranks of group 1 in the competence dimension for the first day and the other half of participants learned the relative ranks of the group 1 in the popularity dimension for the first day.

#### Test blocks of day 1 and day 2: Transitive Inferences

After the learning block, we tested whether participants could generalize their knowledge to infer the relative status between individuals having one or more rank-level differences. This test block followed each learning mini-block (the test blocks in **Fig. S1**). During the test block, all possible pairs of within-group individuals were presented to participants except for the pair of individuals who are at the same rank in the given dimension (meaning there was always a correct answer). In each trial, participants were choosing the superior face in the given dimension. Participants were instructed that their choices would count towards their final payout (the greater the payout when overall accuracy >90%). No feedback was given during test blocks to prevent further learning.

#### Test 2 blocks of day 2: Flexible inferences in intermixed behavioral contexts

At the end of the second day of training, an additional test block was presented. During this test 2 block, trials asking the relative rank of group 1 individuals and group 2 individuals were intermixed, as was the task-relevant dimension (the test 2 block in **Fig S1B**). Otherwise, these trials were identical to the other test blocks.

#### Hub learning block of day 3: Learning ‘hubs’ between-groups

On the third day of training, participants learned the relative ranks of pairs of between-group individuals for the first time. Importantly, the purpose of the hub learning was to provide limited experience about relative ranks of certain between-group individuals. That is, participants did not learn the relative rank of all pairs of between-group individuals but only the relative rank of selected pairs of between-group individuals.

During the hub learning block, participants learned the relative status between one individual in one group (hub) and another individual in the other group (non-hub) with one rank level difference in the given dimension (**Fig. S2A**). In each group, four individuals (two per group) were selected as hubs in one dimension. In the other dimension, a different four individuals played a role as hubs (eight hubs in total; **Fig. S2E**). For those two hubs in each group, one was at the second rank, and the other was at the third rank in the given dimension. Each of the hubs was paired with four different individuals in the other group (See the possible pairs in **S2B** and **S2C**). With this procedure, all eight individuals in one group (non-hub; **Fig S2D**) were paired with two selected individuals in the other group in one dimension (they were never paired with the other six individuals who were not selected as hubs). In particular, a hub in group 1 who was at the third-rank in the dimension was paired with four non-hub individuals in group 2 including two second-rank individuals and two fourth-rank individuals (the top panel in **Fig. S2B**). The other hub in group 1 who was at the second-rank in the given dimension was paired with the other four individuals in group 2 including two first-rank individuals and two third-rank individuals (the bottom panel in **Fig. S2B**). This is also true for hubs in group 2 (**Fig. S2C**). During hub learning, participants have therefore learned the relative rank of some pairs of between-group individuals who have one rank difference, as they learned for the pairs of within-group individuals during the previous learning blocks. The hub learning block allowed us to create a unique “path” between members of different groups. That is, each of 12 non-hubs individuals (six per each group; **Fig. S2D**) has a unique connection to a specific hub in the other group (one among eight hubs in **Fig. S2E**) in one of two hierarchy dimensions. Note that the hubs in competence dimension differed from the hubs in popularity dimension (e.g. **Fig S2F**).

For each trial in the hub learning block, three face stimuli (F1, F2, and F3) were presented for 2 s sequentially after the presentation of a conditional cue (1 s) indicating the task-relevant dimension of the current trial (**Fig. S2A**). Participants were asked to indicate one who is superior rank between F1 and F2 in the given dimension while F2 was presented. Feedback (correct/ incorrect) followed after each decision (2 s). Between F1 and F2, one was the hub in the given dimension and the other was a non-hub in the different group who has one rank difference from the hub. A third face (F3) was presented (2 s) at the end of every trial, and participants were asked to press a button indicating the gender of the F3 face stimuli. F3 was selected from 12 non-hubs in the given dimension (**Fig. S2D**). By presenting non-hub faces at the F3, we controlled the number of presentations of each of the face stimuli to be equivalent. No feedback was given for the gender discrimination task. The inter-stimulus interval (ISI) was 2 s and the inter-trial interval (ITI) was 4 s. While learning between-group relationships via hubs, participants simultaneously became familiar with the task procedure that we used for the fMRI experiment (**Fig. 1F**).

#### Sample size calculation and participant retention

The sample size was determined on the basis of a power calculation using G*Power assuming a medium to large effect (d□= □0.6), and resulting in a sample size of 24 to achieve a statistical power of 80% (α□= □0.05, two-tailed test).

For the behavioral training, we recruited 282 participants. They received course credit as compensation. Among participants who completed the two days of training, 82 participants achieved a higher accuracy than our threshold during the ‘flexible inferences’ block in day 2 (participants who successfully distinguished the second and third rank individuals above chance while also reaching >85% accuracy overall). We included a high-performance threshold because we needed to ensure accurate representations of cognitive maps, should they exist, to be able to measure them reliably. Moreover, the relatively high drop-out rate in part reflects difficulties retaining subjects for three-day studies, variable performance due in part to using course credit as incentives (e.g. many students had achieved full credits for their courses before the end of Day 2 training), and true variability of task performance.

Among 65 participants who agreed to continue on Day 3 training with monetary compensation, 51 participants made correct inferences during the hub learning more than at chance level during the last training session of the day 3 training. Of these 51 participants, 33 further participated in the fMRI experiment and 18 participated in a behavioral version of the inferences task (see **Fig. S5**).

### fMRI experiment

The purpose of the fMRI experiment paradigm was to test whether and how participants represent their knowledge of social hierarchies of the two groups of individuals and make inferences about relative ranks of novel pairs of individuals. **Fig. 1F** illustrates an example trial of the fMRI experiment. In each trial, three face stimuli (F1, F2, and F3) were shown sequentially following a conditional cue (1 s) with an inter-stimuli fixation cross (1.5 s). The color of a square shown as the conditional cue indicated the relevant dimension of the current trial. These conditions were equated and randomly interleaved. Each face stimulus was shown for 2 s. Between face stimuli, we presented a fixation cross for inter-stimuli-intervals (ISI) which were jittered between 2 ∼ 5 pulses (TR=1200ms). The first decision was made during the F2 presentation. Participants were asked to press a button to indicate who is superior rank between F1 and F2 in the given social hierarchy dimension. They were asked to respond as quickly as possible but also as accurately as possible. No feedback was given. The second decision was made during the F3 presentation. Participants were asked to press a button to indicate the gender of F3 as quickly as possible. The buttons allocated to indicate the gender of presenting face stimuli were counterbalanced across participants. If the response was missed in the inference decision due to a slow response, we showed a ‘missed’ sign and proceeded to the next trial. The missed trial was then tested again after a random number of trials, which allowed us to collect responses to all trials from all participants.

The following was *not* told to participants: (1) during the fMRI experiment, F1 and F2 were selected from different groups among 12 non-hubs individuals in the given dimension (**Fig. S2D**) – F1 was selected from group 1 for one-half of trials and F2 was selected from group 1 for the other half; (2) F3 was selected from among eight individuals who played a role as hubs regardless of the social hierarchy dimension (**Fig. S2E**).

All eight hubs were shown the same number of trials at the time of the F3 presentation. Participants were asked to make the same type of decisions as they did for the third-day behavioral training (i.e. choosing a higher rank individual between the first two faces in the given context dimension and indicating the gender of the third face). However, unbeknownst to participants, all pairs were novel (i.e. they had never been compared before). This manipulation meant we were able to test whether and how participants make inferences about the relative rank of unlearned pairs of individuals. The fMRI experiment comprised two blocks. Each block included 104 trials which included all possible between-group pairs of non-hubs who have different ranks in the given dimension. Note that, during the fMRI experiment, all F1-F2 pairs were also presented in reverse order in both context dimensions. The order of the trials was randomized across participants.

#### Inferences of relative status of unexperienced pairs via hubs

During training, participants never directly learned the relative status between two face stimuli (F1 and F2) presented during the fMRI paradigm. Instead, participants could make transitive inferences about relative status of unlearned pairs via one of two hub individuals (H1 and H2), (**Fig. 1D**). Note that, for every F1-F2 pair, there were only two individuals (H1 and H2) that have been paired with either F1 or F2 during training in the given dimension. That is, H1 belonged to the same group with F2 (within-group) which had been uniquely paired with F1 during the hub learning block (between-group). Likewise, H2 belonged to the same group with F1 (within-group) which had been uniquely paired with F2 during the hub learning block (between-group). The direction and the distance of inference trajectories on the social cognitive map were, therefore, determined by which of the hubs (H1 or H2) was preferentially recalled by participants for making inferences. The between-group relationship to the hub (F1 → H1 and F2 → H2) had one rank difference in the given dimension. If participants recalled H1, the transitive inference depended on the within-group distance (H1 → F2), and the inference was made in the forward direction (F1 → F2). If participants recalled H2, the inference depended on the within-group distance (H2 → F1), and the inference was made in the backward direction (F2→ F1). We examined which trajectory participants used for making transitive inferences by examining which unseen hub was selectively retrieved during inferences.

### Behavioral data analysis

We analyzed the reaction times (RT) and accuracy in inferences of the relative status between a novel pair of individuals (F1 and F2). The choice RT was measured from the F2 onset to the response. To make successful inferences, participants could use the cognitive map of social space to make inferences via an unseen hub. The inference trajectories, therefore, were grounded by the location of the hub. To examine whether either or both hubs were selected for inferences, we regressed choice RT on different distance measures of putative inference trajectories in the same multiple linear regression model. We focused our regression analysis on choice RT only because choice accuracy showed a ceiling effect (**Fig. S3B**). As regressors, we included both distances which were measured from each of two potential hubs: the distance between H1 and F2 and the distance between H2 and F1 in addition to the distance between F1 and F2 by allowing them to compete to explain RT variance. Moreover, the distance was measured in both of the rank difference in the task-relevant dimension (D) and Euclidean distance (E). We regressed RT of inference decisions on four different distance measures of trajectories via hubs, *D*_*H1F2*_, *D*_*H2F1*_, *D*_*H1F2*_, and *D*_*H2F1*_(**Fig. 1H**) and two direct distance measures between F1 and F2, *D*_*F1F2*_ and *E*_*F1F2*_(**Fig. 1I**), (**Eq. 1**).

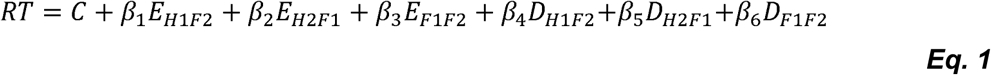

The correlation between different distance measures is shown in **Fig. S6B**. Group level effects of each of the distance measures were tested with a one-sample t-test to account subjects as a random variable.

### Functional imaging acquisition

We acquired T2-weighted functional images on a Siemens Skyra 3 Tesla scanner. We used gradient-echo-planar imaging (EPI) pulse sequence that sets the slice angle of 30° relative to the anterior-posterior commissure line, minimizing the signal loss in the orbitofrontal cortex region (Weiskopf, Hutton, Josephs, & Deichmann, 2006). We acquired 38 slices, 3mm thick with the following parameters: repetition time (TR) = 1200 ms, echo time (TE) = 24 ms, flip angle = 67°, field of view (FoV) = 192mm, voxel size = 3 × 3 × 3 mm3. Contiguous slices were acquired in interleaved order. We also acquired a field map to correct for potential deformations with dual echo-time images covering the whole brain, with the following parameters: TR = 630 ms, TE1 = 10 ms, TE2 = 12.46 ms, flip angle = 40°, FoV = 192mm, voxel size = 3 × 3 × 3 mm3. For accurate registration of the EPIs to the standard space, we acquired a T1-weighted structural image using a magnetization-prepared rapid gradient echo sequence (MPRAGE) with the following parameters: TR = 1800 ms, TE = 2.96 ms, flip angle = 7°, FoV = 256mm, voxel size = 1 × 1 × 1 mm3.

### Pre-processing

The preprocessing of functional imaging data was performed using SPM12 (Wellcome Trust Centre for Neuroimaging). Images were corrected for slice timing, realigned to the first volume, and realigned to correct for motion using a six-parameter rigid body transformation. Inhomogeneities created using the phase of nonEPI gradient echo images at 2 echo times were coregistered with structural maps. Images were then spatially normalized by warping subject-specific images to the reference brain in MNI (Montreal Neurological Institute) coordinates (2mm isotropic voxels). For the univariate analysis images were smoothed using an 8-mm full-width at half maximum Gaussian kernel (Mikl et al., 2008).

### Univariate analysis

We implemented several general linear models (GLMs) to analyze the fMRI data. All GLMs contained separate onset regressors for the contextual cue which indicates the task-relevant dimension, F1, F2, and F3 stimuli presentations for each of the trials. Specifically, the F3 onsets were separately modeled when F3 was (1) the task-relevant hub, H1, (2) the task-relevant hub, H2, and (3) neither H1 nor H2, but the hub for other pairs of individuals (non-relevant hub). A stick function modeled the contextual cue and the F3 presentation and a 2 s boxcar function modeled the presentation of F1 and F2. The F1 onset regressors were modulated with parametric regressors of the rank of the individual in the task-relevant hierarchy (F1_R_) and the rank in the task-irrelevant hierarchy (F1_I_). The F2 onset regressors were modulated by the rank in the task-relevant hierarchy (F2_R_), the rank in the task-irrelevant hierarchy (F2_I_), and additional regressors of interest representing the putative inference trajectories which varied according to which structure of cognitive map was tested (1-D or 2-D) (**Fig. S6**). The onset of button presses (stick function) and the 6 motion parameters obtained during realignment were entered into the GLM as regressors of no interest. The orthogonalization function was turned off. All these regressors, except for the motion parameters, were convolved with the canonical hemodynamic response function in SPM12.

To test whether the brain encodes the trajectories via hubs over Euclidean space, for **GLM1**, we included parametric regressors of Euclidean distance and cosine angle of the vector between F1 and H2 and those of the vector between F2 and H1 (E_H2F1_, A_H2F1_, E_H1F2_, and A_H1F2_). The Euclidean distance between face stimuli was defined over the two-dimensional (2-D) space characterized by their relative rank in each of two social hierarchies. The cosine angle represents the normalized function of competence modulated by popularity. The value of these regressors was invariant to the relevant dimension for the current trial.

From the first-level analysis, contrast images of parameter estimates from regressors of inference trajectories (E_H2F1_, A_H2F1_, E_H1F2_, and A_H1F2_) at the time of F2 presentation were estimated from each of the participants. In addition, during the cover task (at the time of F3 presentation), the following contrasts images were estimated for the cross stimuli suppression analysis: trials when F3 was the task-relevant hub, H1, compared to when F3 was a non-relevant hub (H1 < Non-relevant hub); and when F3 was the task-relevant hub (H2) compared to when F3 was a non-relevant hub (H2 < Non-relevant hub).

For **GLM2**, we tested whether the brain uses different cognitive maps for each dimension learned on a different day (popularity and competence), for which we would predict task-modulated distance terms for current behaviorally-relevant and irrelevant task dimensions. We therefore inputted the rank difference in the task-relevant dimension (D) and that of the task-irrelevant dimension (I) as the parametric regressors, which includes the 1-D distances between H2 and F1 and those of H1 and F2 (D_H2F1_, I_H2F1_, D_H1F2_ and I_H1F2_), in addition to the other regressors not associated with the distance measures that we inputted in GLM1. The value of these regressors was dependent on the task-relevant dimension on the current trial.

For **GLM3**, we tested whether the brain has already integrated the cognitive map for group 1 and that for group 2 into a combined cognitive map and encodes the inference trajectories between F1 and F2. We included the regressors of Euclidean distance and cosine angle of the vector between F1 and F2 (E_F1F2_ and A_F1F2_), in addition to the other regressors that we inputted in GLM1.

For **GLM4**, we tested whether the brain uses different combined cognitive maps for making inferences in different contextual dimensions. We inputted the rank difference in the task-relevant dimension and that of the task-irrelevant dimension as 1-D distances between F1 and F2 (D_F1F2_ and I_F1F2_) in addition to the other regressors that we inputted in GLM1. **Fig. S6A** illustrates the regressors of different models to examine how the brain constructs and use a cognitive map of abstract social hierarchies.

### Group-Level Statistical Inference

We perform group-level inference both on hypothesis-driven *a priori* ROIs in the HC, EC, and vmPFC/mOFC bilaterally and exploratory whole-brain analyses. For our *a priori* ROIs, we reported our results in these areas at the threshold *p*_*TFCE*_<0.05 using threshold-free cluster enhancement (TFCE) (Smith & Nichols, 2009) for correction of multiple comparisons within a combined ROI which integrated anatomically defined HC, EC, and vmPFC/mOFC in bilateral hemispheres into a single mask. For the whole brain analyses, we enter the individual contrast images into the second-level analysis. For the whole-brain analysis, we reported the whole-brain permutation-based threshold-free cluster enhancement (TFCE)-corrected images at the threshold *p*_*TFCE*_<0.05 (1000 iterations of simulation).

### Repetition suppression analysis

Recent findings have shown that blood-oxygen-level-dependent (BOLD) suppression can be measured not only to repetition of a stimulus, but also to pairs of stimuli that have been well-learned though association and to a predicted or imagined outcome (Barron et al., 2016; Boorman et al., 2016; Grill-Spector et al., 2006; Klein-Flugge et al., 2013). This cross-stimulus suppression (CSS) approach allows us to examine the underlying neural representations of retrieved memories. In the current study, if the relevant hub is presented during the suppression phase, at the time of F3 presentation, directly after participants recall the relevant hub for making inferences of relative ranks between faces, then the BOLD signal in the areas reinstating the relevant hub should be suppressed compared to the non-relevant hubs. Considering that effects of CSS did not depend on the responses of participants during the cover task, we included the BOLD responses in every F3 presentation into the analysis. Moreover, the BOLD signal should be suppressed specifically for the relevant and preferentially selected hub compared to the relevant but unselected hubs. Considering that the hippocampus (HC) was our *a priori* ROI, we reported our results at a threshold p_TFCE_<0.05 using TFCE within a anatomically defined independent ROI that includes bilateral HC for correction of multiple comparisons.

It is important to note that the hub face in each trial was not determined by F3 stimuli but by the combination of task-relevant dimensions, F1 and F2. That is, the same face stimuli presented at F3 could be the relevant hub (e.g. H2) in one trial but it could be an alternative hub (e.g. H1) or non-relevant hub (the faces that have not been paired with both F1 and F2 in the given dimension) in other trials. Therefore, the effects of RS were not driven by the comparison of specific stimuli set but the comparison between conditions.

### Neural model comparison

The different hypothetical structure of the cognitive map and putative inference trajectories cannot be accurately tested in a single GLM while there is potential multicollinearity between different distance measures. By definition, this was the case for some of our distance terms because, for example, E is correlated with D and I. The cross-correlation between different distance measures is shown in **Fig. S6B**. To formally compare the predictability of each distance measure in different models, we therefore used Bayesian model selection (BMS), (Stephan, Penny, Daunizeau, Moran, & Friston, 2009).

We tested whether neural activity in the vmPFC/mOFC and EC, which showed effects of the Euclidean distance of the inference trajectory from the hub (*E*_*H2 F1*_), is better explained with alternative distance measures of inference trajectories. To test this, we ran several GLMs in which the brain activity at the time of inferences (F2 presentation) was modeled with only one of candidate distance measures as a parametric regressor. Specifically, we compared the models having one of the following distance measures as parametric regressors: *E*_*H2F1*_, *E*_*H1F2*_, *E*_*F1F2*_, *D*_*H2F1*_, *D*_*H1F2*_, *D*_*F1F2*_, *A*_*H2F1*_, *A*_*H1F2*_, and *A*_*F1F2*_. The inference process can alternatively be modeled with the link distance (*L*), which indicates the minimum number of links between F1 and F2 in the social network. The shortest link distance, *L* equals the sum of the number of links from F2 to H2 (between-group) and the number of links from H2 to F1 (within-group). Since we controlled the between-group distances as one, the brain areas encoding *L* can be estimated by the GLM which includes *D*_*H2F1*_ as a parametric regressor. In addition to the parametric regressors of distance, all GLMs also included the rank of the task-relevant dimension and the rank of the task-irrelevant dimension of presenting faces as additional regressors at the time of F1 and F2 presentation (F1_R_, F1_I_, F2_R_, and F2_I_). As before, the onsets of the contextual dimension cue, F3 presentation, and button presses were also entered as additional regressors in all GLMs.

For the univariate neural model comparisons, we first estimated the log-likelihood of each of GLMs. Following previous work (Kumaran et al., 2016; Niv et al., 2015; Wilson & Niv, 2015), the log-likelihood (LL) of each of the models was calculated (**Eq**.**2**) separately for the *a priori* anatomically defined ROIs: the EC and vmPFC/mOFC.

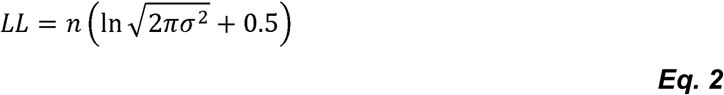

where *n* is the total number of scans, and σ^2^ is the variance of the residuals after subtracting the best-fit linear model. Considering that the linear model provides the maximum likelihood solution to each model with Gaussian-distributed noise, the likelihood was computed from residuals in the ROIs after subtracting the best-fit linear model. Since all models had the same number of parameters, their likelihoods could be directly compared to ask which model accounted best for the neural activation patterns. We further entered the LL to Bayesian model selection (BMS) to compare the goodness of fit of the model with the exceedance probability (XP).

### Regions of Interest (ROI) analyses

The ROIs were defined in the bilateral HC (Yushkevich et al., 2015), bilateral EC (Amunts et al., 2005; Zilles & Amunts, 2010), bilateral amygdala (AM) (Tzourio-Mazoyer et al., 2002), and bilateral vmPFC/mOFC (F.-X. Neubert et al., 2015) using probabilistic map of anatomical ROIs. We also included additional ROIs in the bilateral primary motor cortex (M1) (Glasser et al., 2016) as control regions. The ROIs in the HC, AM, and M1 have been already binarized by the authors of each study ((Yushkevich et al., 2015) for HC; (Tzourio-Mazoyer et al., 2002) for AM; (Glasser et al., 2016) for M1). Neubert et al. also provided the binarized ROIs in vmPFC/mOFC (F.-X. Neubert et al., 2015) and noted that they set the maximum threshold to 25 meaning it includes voxels that belong to any given mask in 25-100% of participants. The EC defined by Juelich atlas (Amunts et al., 2005; Zilles & Amunts, 2010) allowed us to choose the threshold of images based on probability. To define ROIs in the EC, we binarized the probabilistic map in which the minimum threshold was 0 and the maximum threshold was 10. Note that ROIs were independently defined from the current task. For display purposes, all statistical parametric maps presented in the manuscript are unmasked.

### ROI-based representational similarity analysis (RSA)

#### Neural representation of social hierarchies

In hypothesis-driven analyses, we performed a representational similarity analysis (RSA) (Kriegeskorte, Mur, & Bandettini, 2008; Nili et al., 2014) to test whether the *a priori* ROIs contain hypothesized cognitive map structures with respect to the social hierarchies. To test our hypotheses, we first estimated Beta coefficients when each of the individual faces was shown at the time of F1 or F2 in each of our same anatomical ROIs in HC, EC, and vmPFC/mOFC. We then averaged the unsmoothed Beta maps across F1 and F2 presentations, allowing us to estimate the patterns of neural activity in each ROI. These patterns were separately estimated according to which social hierarchy dimension (competence or popularity) was relevant to the current decision. Reliability of data was improved by applying multivariate noise normalization (Walther et al., 2016). We quantified the representational similarity for the two independent fMRI blocks (i.e. runs) using the Mahalanobis distance between the activity patterns, which generated a 24×24 representational dissimilarity matrix (RDM; 12 non-hub individuals were presented in each of two dimensions; **Fig. 4A**). These analysis steps were repeated per ROI.

Next, we confirmed that the RDM estimated from the brain activity patterns in each of the ROIs discriminated different face stimuli with good sensitivity using the exemplar discriminability index (EDI) (Nili, Walther, Alink, & Kriegeskorte, 2016), which is defined as the average of the pattern dissimilarity estimates between different stimuli compared to the average of the pattern dissimilarity estimates between the same stimuli. We confirm that the EDI in all ROIs was positive (one-sample t-test, p<0.01) suggesting that the different sets of stimuli were discriminable based on their multivariate activity patterns.

We predicted the RDM estimated from the patterns of neural activity in *a priori* ROIs using several candidate model-based predictor RDMs (Model RDM; **Fig. 4B**). The model RDMs included (1) pairwise Euclidean distances between individuals in 2-D social space (*E*); (2) pairwise rank difference between individuals in the task-relevant hierarchy (*D*); (3) the context of which social hierarchy dimension (competence or popularity) the face stimulus was presented in (*C*, task-relevant dimension); (4) which group the face stimulus belonged to during training (*G*).

The extent to which the brain RDM of each ROI was explained by the model RDM was estimated with the rank correlation (Kendall’s *τ*_A_). For group-level inference, this effect was then tested at the group-level with the non-parametric Wilcoxon signed-rank test across participants (Nili et al., 2014). The ROI-based analysis uses the pattern of beta coefficients across voxels in the entire ROI. Therefore, correction for multiple comparisons was made over the number of ROIs (n=6), as well as by the number of model RDM comparisons (n=4). We report results corrected for family-wise error (FWE) with the Holm-Bonferroni method at p<0.05, but also note stronger effects with asterisks.

#### Partial correlation analyses

In addition to the model RDMs, using partial correlation, we also tested for an effect of E while controlling for the shared covariance between E and D. Specifically, we estimated the extent to which the RDM estimated in each ROI was explained by E’. E’ denotes the pairwise Euclidean distances between individuals (E) while regressing out its partial correlation with the pairwise rank difference in the task-relevant distance (D): *E′* = *E* − *DD*^+^*E* where *D*^+^ is the Moore-Penrose generalized matrix inverse (*D*^*+*^ *= pinv*(*D*)), **Fig. S8B**). This partial correlation method gives an advantage over the other methods such as orthogonalization which often lose the original correlation structure (**Fig. S8B**). Note that, as **Fig. S8B** shows, E’ differs from E^Orth^ which denotes E orthogonalized by D using the Gram-Schmidt method, or the pairwise rank difference in the task-irrelevant dimension (I). After regressing out the partial correlation, the predictors are independent from each other while preserving high correlation with its original correlation structure. That is, E’ does not correlate with D while it still highly correlates with E (right panel in **Fig**.**S8B**).

Last, we examined the relationship between pattern dissimilarities in each of the ROIs and pairwise Euclidean distances (E), the pairwise task-relevant rank differences (D), and the pairwise task-irrelevant rank differences (I) between all individual faces in the social cognitive map. The brain RDM of each participant was normalized into a range between 0 and 1. The upper triangular part of the normalized 24×24 RDM was arranged according to the distance measure in each model RDM, providing model predictions of representational dissimilarity. We estimated the mean pattern dissimilarity per bin across participants. This analysis was only performed for visualization purposes (**Fig. 4D** for E, **Fig. 4E** for D, and **Fig. 4F** for I) Using the same methods, we also showed the effects of pairwise Euclidean distance (E) between faces and the pattern dissimilarity which were separately analyzed for within-group (E_wth; **Fig. S9A**) and between-group relationships considering the group effects (G). The between-group relationships were separately analyzed also based on whether the faces had been directly compared during training (E_btw_hub; **Fig. S9B**) or not (E_btw_non; **Fig. S9C**). We did not make any statistical interpretation based on this analysis.

### Searchlight-based RSA

Whole-brain searchlight RSA was performed to examine brain areas in which the activity patterns reflect the hypothesized relational structure of the social hierarchies both within and outside of HP, EC, and vmPFC/OFC. Moreover, the searchlight analysis allows us to examine to what extent the model RDM explains the neural representation dissimilarity with a fixed number of voxels examined across regions. We defined a sphere containing 100 voxels around each searchlight center voxel. Consistent with the ROI analysis, we estimated the neural activity patterns elicited while each of the individuals was presented at the time of F1 or F2 from each of the searchlights. These neural representations were separately estimated according to which social dimension was relevant to the current task. The dissimilarity matrices were quantified with the Euclidean distance between neural patterns estimated from different blocks. For each searchlight, therefore, a 24×24 dissimilarity matrix was generated based on neural activity patterns elicited by each of face stimuli in two different task-relevant dimensions (**Fig. 4A**). We used the same predictors (i.e. model RDMs) that we used for ROI-based RSA analysis to estimate the neural representational dissimilarity across searchlights with Kendall’s *τ*_A_ rank correlation. To assess neural dissimilarity specific to E, we also tested the RDM E’ while partialling out its covariance with D. The computed Kendall’s *τ*_A_ values were then mapped back on the central voxel, allowing continuous mapping of information in the whole-brain per subject. These images were further smoothed using an 8-mm full-width at half maximum (FWHM) Gaussian kernel and Fisher’s Z transformed. We further performed one-sample t-tests for a group-level analysis. We corrected for multiple comparisons using TFCE (Smith & Nichols, 2009) with 1000 iterations of simulation. We reported the results corrected for family-wise error (FWE) for multiple comparisons across searchlights (p_TFCE_<0.05).

