## Supplementary material for "Map making: Constructing, combining, and inferring on abstract cognitive maps": Fig. S

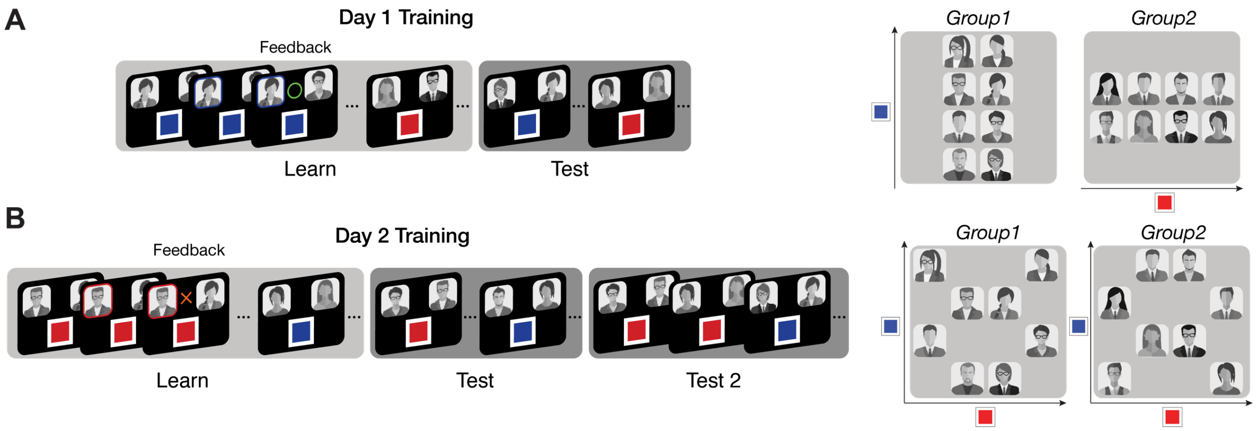


**Figure S1.** Behavioral training for day 1 and day 2. **A.** During the Learn phase on day 1, participants learned the relative rank of members in each group in one of two dimensions based on feedback from binary comparisons. They were asked to choose the higher rank individual between two members in the same group who differed by one level only in the given social hierarchy dimension. During the test phase on day 1, participants were asked to infer the relationship between two in the same group who were never paired during training through transitive inferences. No feedback was given during test phase. After day 1 training, participants could have built a hierarchical structure of each of the two groups in one dimension (Right panel). **B.** During the learn phase on day 2, participants learned the relative status of members in each group in the unlearned second dimension by comparing two members in the same group who differed by one level only in the corresponding dimension. During the test phase on day 2, participants were asked to infer the relative status of unpaired individuals through transitive inference. No feedback was given during test phase. At the end of day 2 training, knowledge about within-group social hierarchies in both dimensions was tested (Test 2). During the Test 2 phase, participants were asked to infer the relative status of two individuals in the same group while both groups and dimensions were intermixed across trials. After training on day 2, participants could in principle have built a hierarchical structure of each of two groups in two dimensions (Right panel).

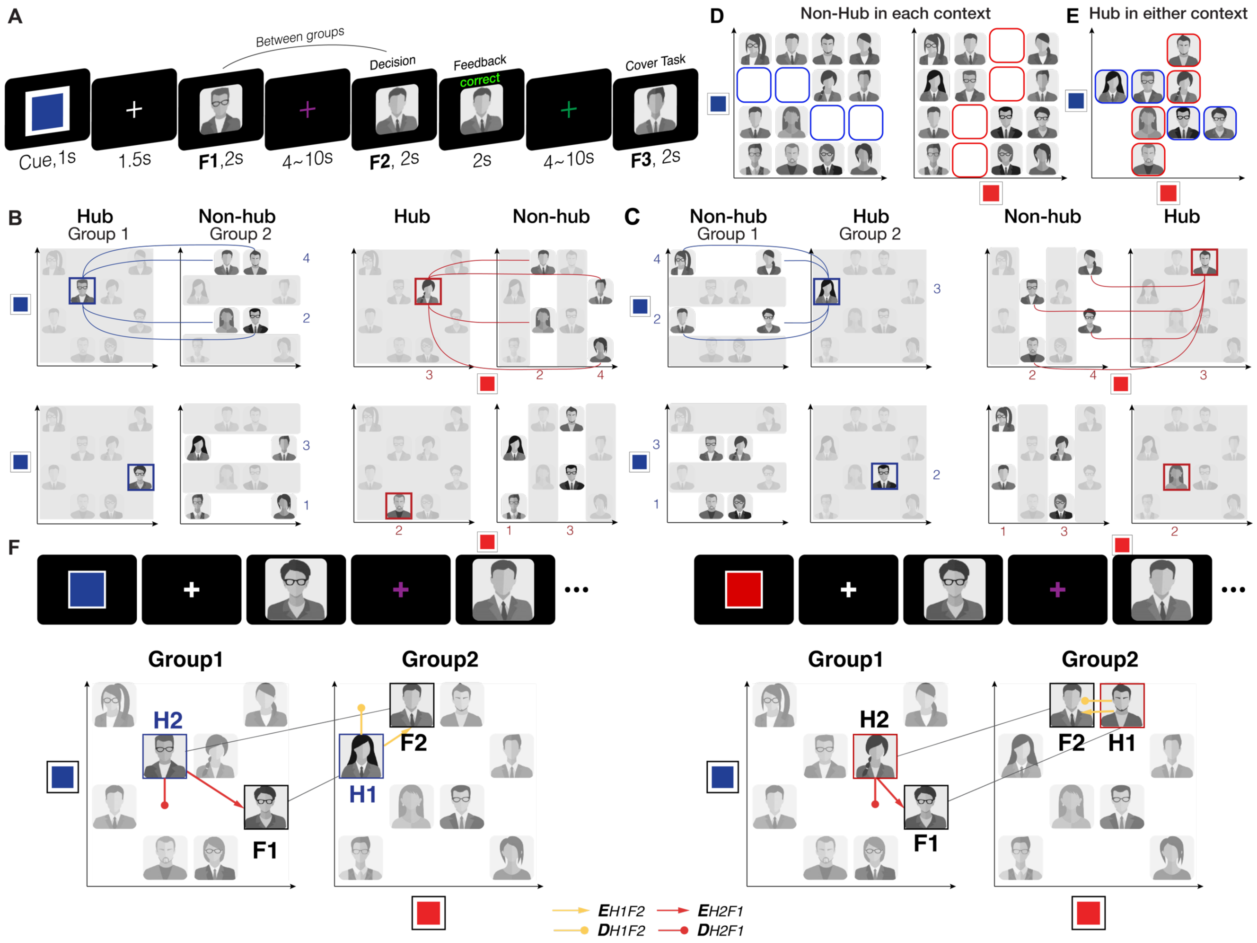


**Figure S2.** Behavioral training on day 3, performed before fMRI scanning on the same day. **A.** Participants made inferences about the hierarchical relationship of two between-group individuals (F1 and F2) in a given dimension (indicated by cue color). A cover task (indicating the gender of the face stimuli, F3) followed at the end of every trial. **B (*C*).** F1 and F2 pairs were selected as follows. In Group 2 (*Group 1*), four individuals whose rank are the 1st or the 3rd in the given dimension are paired specifically to a face stimulus in the other group, Group 1 (*Group 2*), whose rank is the 2nd in the given dimension. The remaining four individuals in Group 2 (*Group 1*) whose rank are the 2nd or the 4th are specifically paired with another member in the other group, Group 1 (*Group 2*), whose rank is the 3rd in the dimension. This is also true in the other dimension (Right panels). We called the individuals in Group 1 (*Group 2*) who had been paired with four other individuals in Group 2 (*Group 1*) 'hubs'. For each trial of the hub learning phase, therefore, participants were asked to make a binary decision comparing between-group individuals including one hub individual who differed by one level on the given dimension. **D.** In each dimension, twelve individuals play a role of 'non-hub'. In fMRI, participants were asked to infer the relative status between non-hub individuals in different groups who had not been directly paired during training. The left panel shows individuals who were shown in the popularity dimension, and the right panel shows individuals who were shown in the competence dimension. **E.** Eight individuals play a role as hubs. Among four hubs in each group, two hubs were for the competence dimension (highlighted in red); two hubs were for the popularity dimension (highlighted in blue). **F.** Importantly, hubs in one dimension differ from those in the other dimension, which means that to make an accurate inference of the relative status of the same pair of individuals in the two different dimensions during the fMRI task, participants needed to retrieve different hubs, which would alter the inference trajectories (e.g. inferring relative status of the same pair individuals, F1-F2 in popularity dimension on left and competence dimension on right).

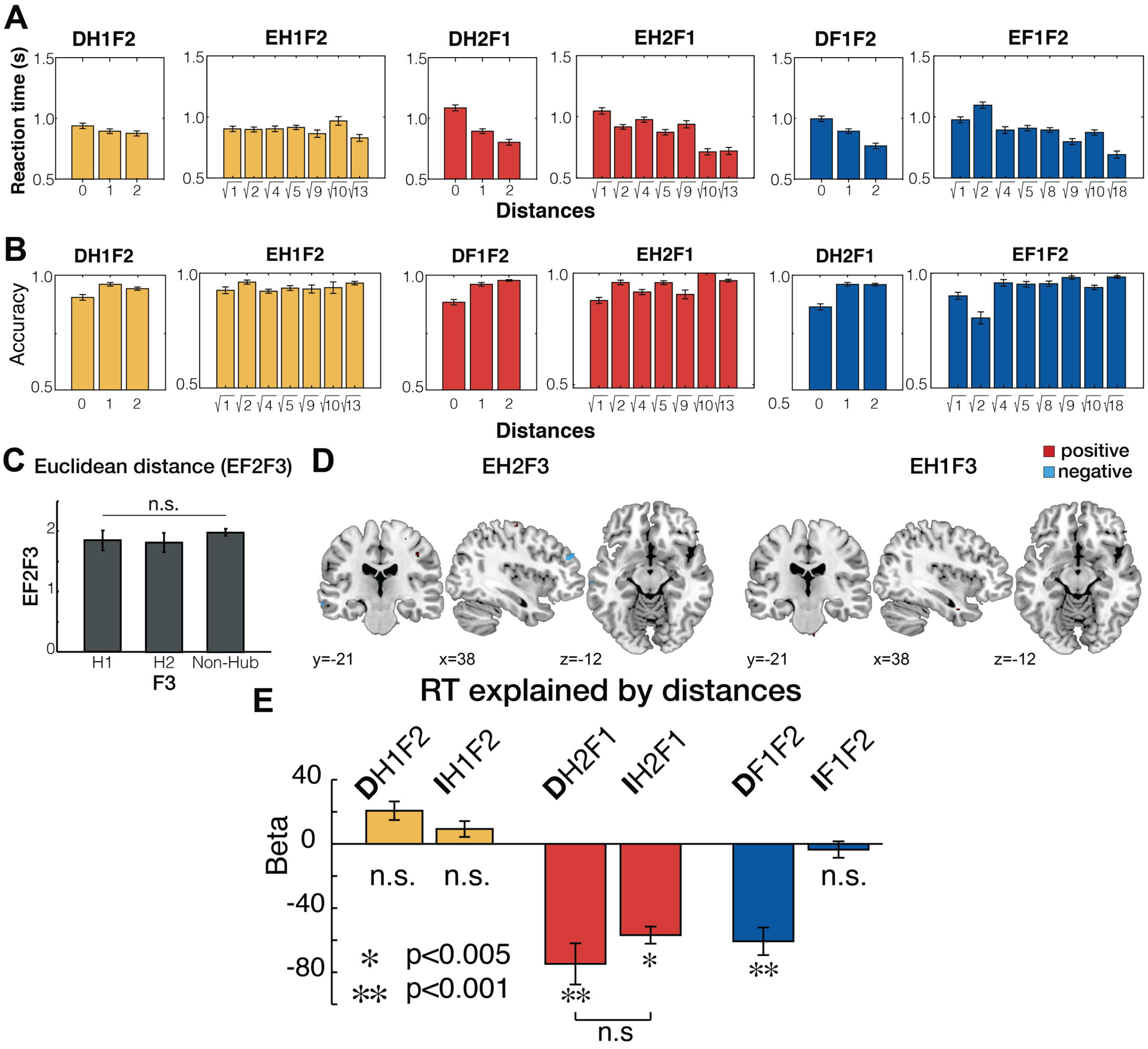


**Figure S3. A.** Changes in reaction time (RT) in inferences as a function of distances of different types of inference trajectories. Note these RT plots do not control for the alternative distance metrics, as the regression analyses do. **B.** Changes in accuracy (% correct) in inferences made in fMRI experiments as a function of distances of different types of inference trajectories. **C.** The face stimuli of F3 were selected from hubs (H1, H2, and "non-hubs"), which controlled for the Euclidean distance from F2 ($E_{F2F3}$). This ensured we could examine the cross-stimulus suppression effect for the relevant hub independently from the relative relationship between F2 and F3 ($F_2$=0.77, p=0.47, One-way ANOVA). **D.** Neural correlates of Euclidean distance from the potential latent hubs (H2 and H1) and F3 ($E_{H2F3}$ and $E_{H1F3}$) at the time of F3 presentation (the brain areas showing a positive correlation are colored in red, and those showing an inverse correlation in blue, p<0.005, uncorrected). We did not find any effects in bilateral hippocampus (HC) even at a lenient threshold, p<0.01, uncorrected. **E**. The Euclidean distance (E) is factorized with two orthogonal vectors, the rank distance in the relevant dimension (D) and the rank distance in irrelevant dimension (I). In addition to the results shown in **Fig. 1J**, we performed an alternative multiple linear regression in which we entered D and I as regressors instead of E. The results showed that both 1-D distances from H2 ($D_{H2F1}$ and $I_{H2F1}$), but not the distances from H1, significantly explain variance in RT, in addition to the direct distance between F1 and F2 ($D_{F1F2}$).

We also found that the effect of $D_{H2F1}$ was not different from the effect of in $I_{H2F1}$ (paired t-test, $t_{26}$=-1.64, p=0.11). Taken together, these results consistently support an effect of $E_{H2F1}$ on RTs, as shown in **Fig. 1J**.

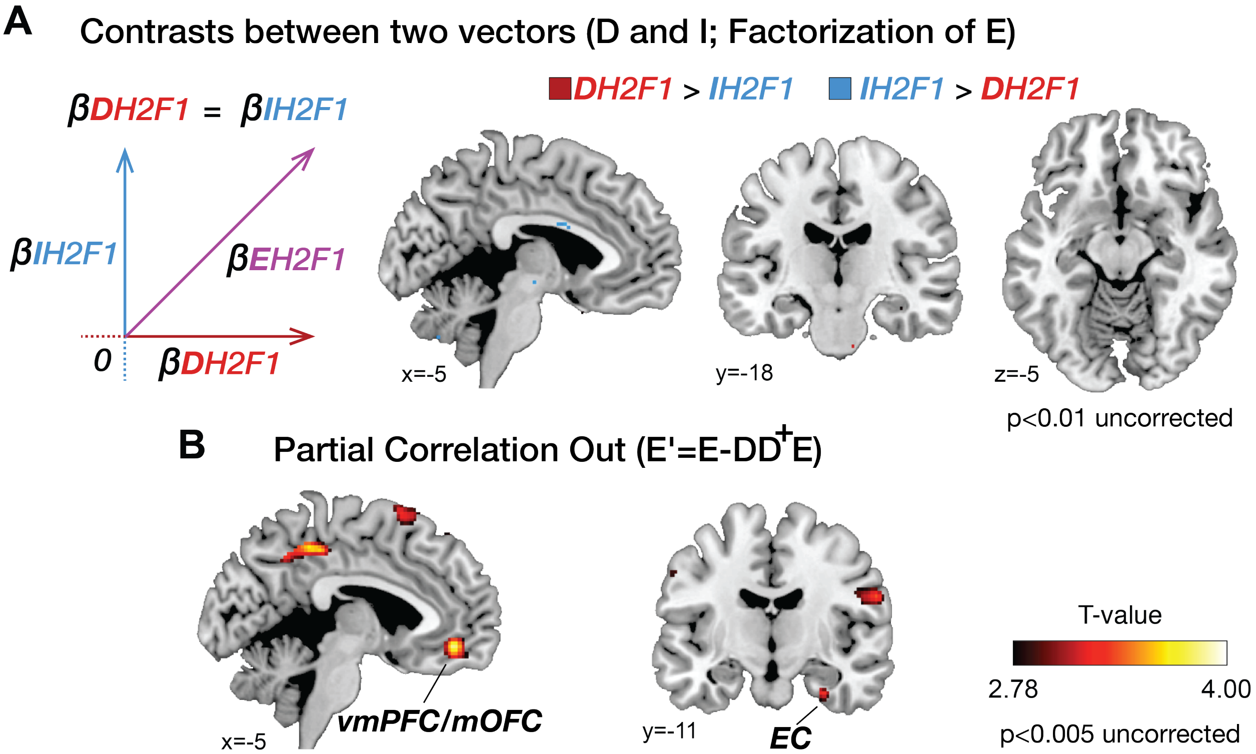


**Figure S4**. **A**. The contrast analysis between positive effects of $D_{H2F1}$ and positive effects of $I_{H2F1}$. This reveals that no brain area preferentially encoded $D_{H2F1}$ or $I_{H2F1}$ over the other even at a liberal threshold (p>0.01, uncorrected). These findings support the conclusion that the brain areas revealed in the conjunction analysis (vmPFC/mOFC and EC in **Fig. 2C**) encode both $D_{H2F1}$ or $I_{H2F1}$ with similar weights, consistent with the interpretation that the vmPFC/mOFC and EC encode Euclidean distance ($E_{H2F1}$). **B**. The effect of $E'_{H2F1}$ which denotes $E_{H2F1}$ after partialling out the1-D task-relevant distance, $D_{H2F1}$: $E' = E - DD^{+}E$, where $D^{+}$ is the Moore-Penrose generalized matrix inverse $(D^{+} = pinv(D))$. We found the effects of $E'_{H2F1}$ in vmPFC/mOFC ([x,y,z]=[6,42,-14], $t_{26}$=3.75, and [x,y,z]=[-12,24,-20], $t_{26}$=3.72) and EC ([x,y,z]=[30,-14,-30], $t_{26}$=3.35) ($p_{TFCE}$<0.05). For visualization purposes, the whole-brain maps are thresholded at p<0.005 uncorrected.

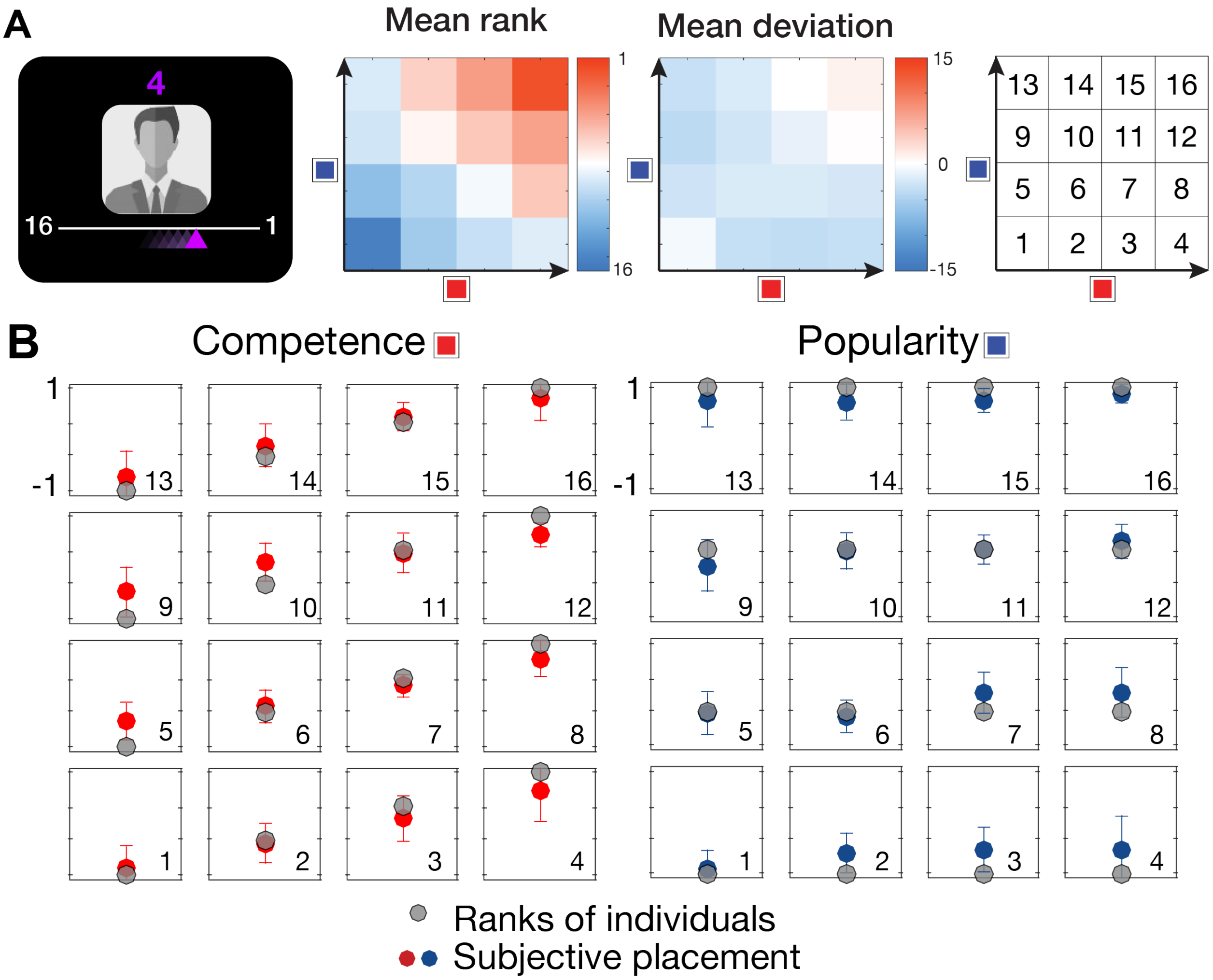


**Figure S5**. A separate group of subjects (n=18) who did not participate in the fMRI part of the experiment performed alternative tasks on day 3, after learning between-group relationships through hubs. Alternative tasks were designed to investigate whether participants can construct a combined cognitive map of 16 individuals (two eight-member groups) according to their ranks in two hierarchy dimensions. **A**. First, participants were asked to rank the 16 individuals according to their 'growth potential (GP)'. We gave the instruction that to compute GP accurately, participants need to weight the ranks in the two dimensions equally. Each of the individuals was presented three times in random order, and participants indicated their rank by moving the cursor on the screen without any time limit (left panel). Mean reported rank (middle panel) and the mean deviation (reported rank - actual rank in GP; right panel) are shown. Participants were able to integrate values from the two dimensions into a single integrated rank value. **B**. Second, participants were asked to place each individual in a 2-D plane where the vertical axis represents the competence dimension and the horizontal axis represents the popularity dimension. 16 face stimuli were shown in a random position. Participants were asked drag-and-drop each of the face stimuli to place them in another position according to their ranks. Responses of each participant were normalized in a range from -1 to 1 by maximum vertical and horizontal distances. The red and blue colored dots indicate the mean position (± s.e.m) of each face stimulus, and the grey dots represent their correct position.

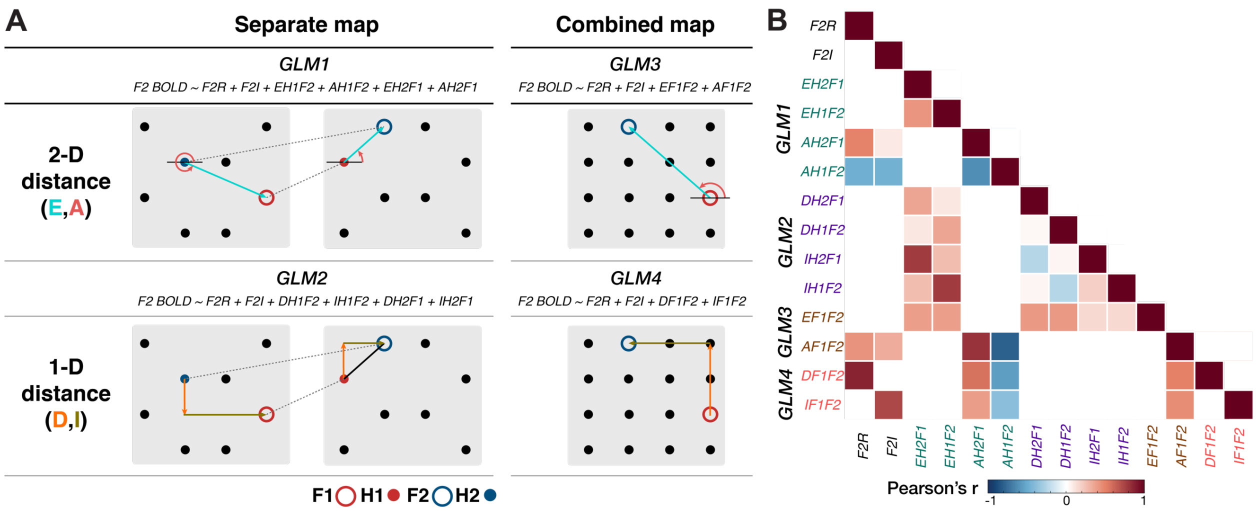


**Figure S6. A.** Four general linear models (GLMs) are depicted to examine the structure the brain constructs to represent social hierarchies and uses to make an inference of relative ranks between F1 and F2. GLM1 tests whether the brain constructs a separate map for each of the two groups and encodes the Euclidean distance from the hub (E) and vector angles between the hub and the connected face (A). GLM2 tests whether the brain constructs a separate map of each group and encodes the one-dimensional (1-D) rank distance in task-relevant dimension (D) and 1-D rank distance in task-irrelevant dimension (I). GLM3 tests whether the brain constructs a combined map and encodes the Euclidean distance (E) and vector angle (A) between F1 and F2. GLM4 tests whether the brain constructs a combined map and encodes D and I between F1 and F2. The task-relevant rank of F2 (F2R) and the task-irrelevant rank (F2I) were also included to model the BOLD signals at the time of F2 presentation, in addition to other common regressors (See Methods). **B.** The cross-correlation (Pearson's r) between different distance metrics for each GLM. In this study, compared to the distance in task-relevant dimension ($D_{H2F1}$ and $D_{H1F2}$), the distances in the task-irrelevant dimension from the hubs ($I_{H2F1}$ and $I_{H1F2}$) had a greater correlation with the Euclidean distances from hubs ($E_{H2F1}$ and $E_{H1F2}$). This was because the distances in the task-irrelevant dimension had a larger variance (I; in a range of 0 to 3) than the distances in the task-relevant dimension from the hubs (D; in a range of 0 to 2), owing to the requirement that hubs were positioned at either rank 2 or 3 in the task-relevant dimension (**Fig. S2 B and C**).

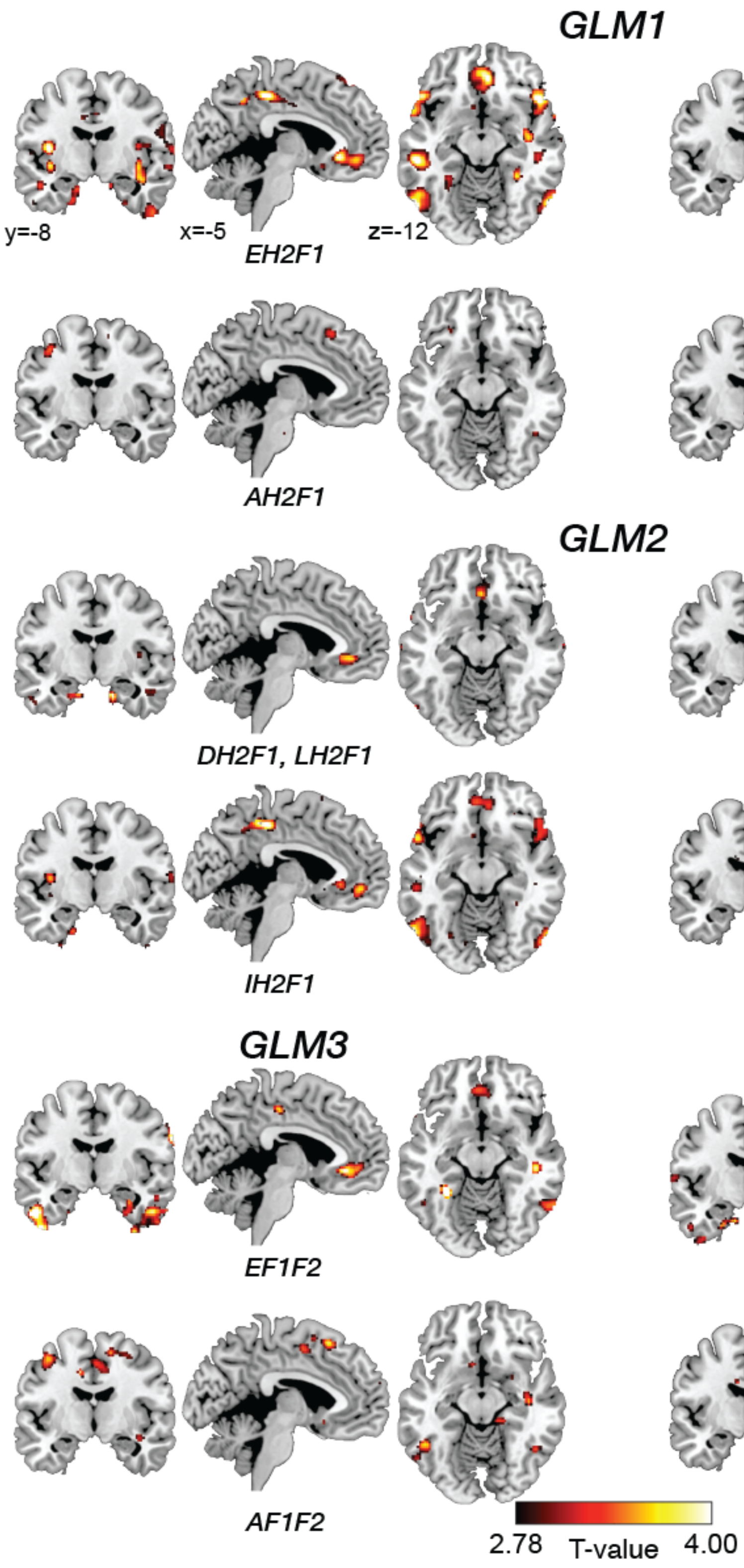


**Figure S7.** In association with **Fig. 2.** Whole-brain univariate parametric analyses showing neural correlates of each of the distance metrics that could have theoretically driven inferences between novel pairs of individuals at the time of decision-making (F2 presentation). Each general linear model (GLM) and inputted regressors are shown in **Fig. S3**. We do note that there was modest evidence that HC activity reflected the vector angle $A_{F1F2}$ (peak [x,y,z,]=[38,-12,-16], $t_{26}=3.56$, $p<0.001$ uncorrected; this effect did not survive at the threshold, $p_{TFCE}<0.05$ in an *a priori* HC ROI), consistent with a previous report (Tavares et al., 2015).

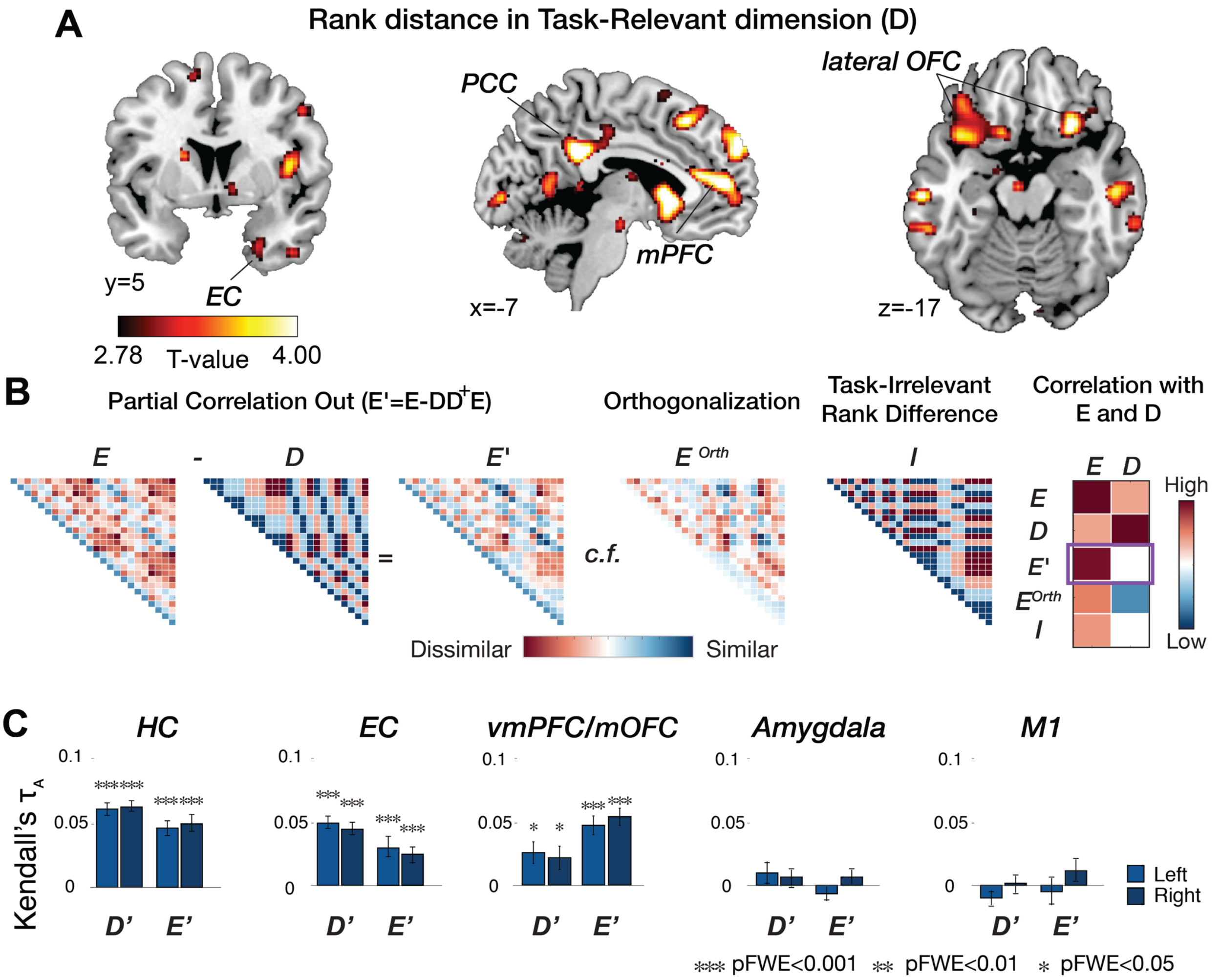


**Figure S8.** In association with **Fig. 4E** and **H. A.** The dissimilarity in the neural activity pattern is explained by model predictions of dissimilarity matrices (model RDM), including pairwise differences in the rank in the task-relevant dimension (D) and pairwise Euclidean distance on the 2-dimension space (E; **Fig. 4G**). Whole-brain searchlight RSA shows effects of 1-D distance in the task-relevant dimension (D) in the EC, lateral OFC, medial prefrontal cortex (mPFC), and posterior cingulate cortex (PCC) ($p_{TFCE}<0.05$). For visualization purposes, the whole-brain maps are thresholded at p<0.005 uncorrected. **B.** We also tested if the effects were specific to each of the model RDMs (D and E) by regressing out their covariance with the other. To do this we subtracted their partial correlation ($E' = E - DD^{+}E,$ where $D^{+}$ is the Moore-Penrose generalized matrix inverse ($D^{+} = pinv(D)$). Specifically, E' was computed as E after regressing out its partial correlation with D. Importantly, E' highly correlates with E but not with D anymore. This partial correlation has an advantage over other methods, such as orthogonalization: E orthogonalized by D ($E^{Orth}$) which creates a negative correlation with D. Moreover, E' differs from the model RDM created by the task-irrelevant rank differences (I). **C.** Representational similarity analysis (RSA) in a priori regions of interests (ROI) including the bilateral HC (Yushkevich et al., 2015), EC (Amunts et al., 2005; Zilles & Amunts, 2010), and vmPFC/mOFC (Neubert, Mars, Sallet, & Rushworth, 2015). The pattern dissimilarity in the brain activity estimated in *a priori* ROIs in increase with E': pairwise Euclidean distance between individuals after partialling out its covariance with D (***, $p_{FWE}<0.001$; **, $p_{FWE}<0.01$; *, $p_{FWE}<0.05$). Conversely, the pattern dissimilarity estimated in the amygdala and primary motor cortex (M1) was neither explained by D' nor E'.

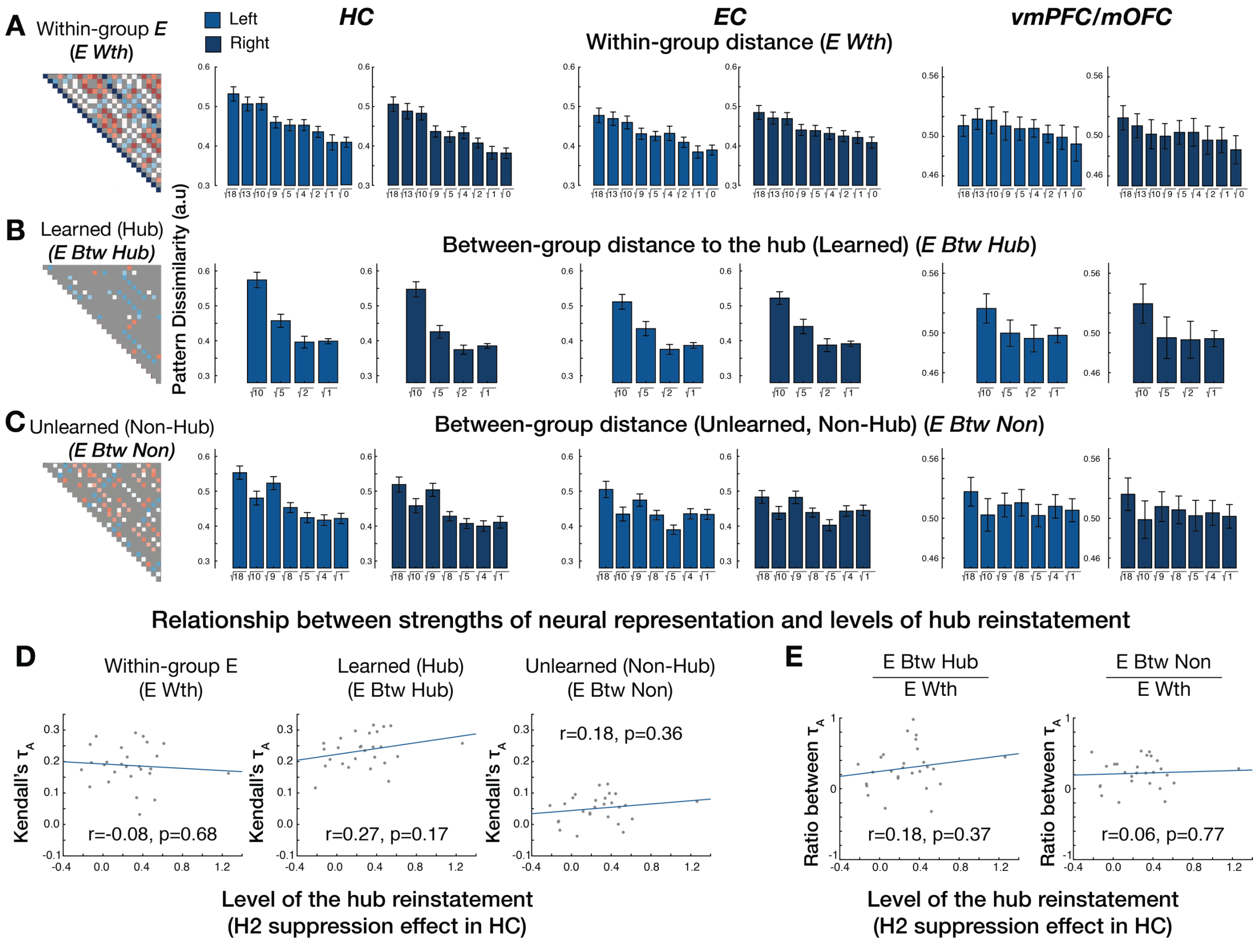


**Figure S9.** In association with **Fig. 4I**. **A.** The dissimilarity between activity patterns estimated in bilateral HC, EC, and vmPFC/mOFC increases in proportion to the pairwise Euclidean distance between within-group individuals (E_wtn). **B**. This is also true for the between-group pairs involving hubs (E_btw_hub). **C**. Compared to those learned relationships, the dissimilarity between activity patterns estimated between non-hub individuals was explained less strongly by E (E_btw_non), although there is a significant effect in hippocampus (HC). 'E wth': Within-group E; 'E btw hub': Between-group E of hub individuals; 'E btw non': Between-group E between non-hub individuals; **D, E.** We tested the possibility that some subjects had fully integrated the two groups into a single map, and therefore, did not need to rely on reinstatement of the hub. Specifically, we conducted an analysis to examine whether individual differences in the level of integration of two social hierarchies correlated with the different levels of reinstatement of the task-relevant hub. The degree of hub reinstatement (x-axis in scatterplots) was estimated by the level of suppression in right HC for H2 compared to non-relevant hubs (**Fig. 3**). To index the strength of social hierarchy representation across participants we used rank correlation (Kendall's $\tau_A$) estimated in the right HC for each type of learned and unlearned comparison (y-axis in scatterplots). $\tau_A$ indicates to what extent the neural activity patterns in the right HC reflect the Euclidean distance of within-group (1), learned between-group (hub) (2), and unlearned between-group (non-hub) (3). **D.** Scatterplot shows the relationship between Hub 2 reinstatement and Kendall's $\tau_A$. With those measures, we found that the level of neural representation explained by pairwise Euclidean distance ($\tau_A$) did not correlate significantly with the level of hub reinstatement across participants. **E.** We also tested whether the level of hub reinstatement was related to the relative strength of the neural representation between (i) unlearned between-group relationships including non-hubs ($\tau_A$ of *E Btw Non*) and learned within-group relationships ($\tau_A$ of *E Wth*) (left panel) and (ii) the relative strength of the neural representation for unlearned between-group relationships including non-hubs ($\tau_A$ of *E Btw Non*) and learned between-group relationships including hubs ($\tau_A$ of *E Btw Hub*) (right panel). We found that these measures of relative strength of neural representation did not correlate significantly with the level of hub reinstatement across participants. This pattern of findings speaks to why participants needed to reinstate the task-relevant hub to make novel inferences before having a representation of a fully combined hierarchy.

**A.**

| | Cluster size | T | Z | Peak coordinate (MNI) x | y | z |
|---|---|---|---|---|---|---|
| | | | EH2F1 | | | |
| right supramarginal gyrus / temporoparietal junction | 2500 | 7.23 | 5.31 | 62 | -24 | 28 |
| right dorsolateral prefrontal cortex / superior frontal sulcus | 35 | 6.27 | 4.85 | 24 | 56 | 30 |
| left supramarginal gyrus / temporoparietal junction | 2451 | 6.02 | 4.72 | -62 | -30 | 34 |
| ventromedial prefrontal cortex / medial orbitofrontal cortex | 443 | 4.82 | 4.04 | 2 | 32 | -6 |
| right posterior middle temporal gyrus | 429 | 4.77 | 4.01 | 62 | -22 | -22 |
| left insula | 62 | 4.53 | 3.86 | -38 | -6 | 10 |
| right insula | 765 | 4.43 | 3.79 | 36 | 0 | 8 |
| right lateral orbitofrontal cortex | 42 | 4.12 | 3.58 | 30 | 34 | -18 |
| right parahippocampal cortex | 14 | 4.04 | 3.53 | 22 | -20 | -26 |
| right superior temporal gyrus | 31 | 3.61 | 3.22 | 56 | -32 | 6 |
| left posterior middle temporal gyrus | 14 | 3.21 | 2.92 | -60 | -52 | -20 |

**B.**

| | | | | | | |
|---|---|---|---|---|---|---|
| | | | DH2F1 | | | |
| left supramarginal gyrus / tempoparietal junction | 968 | 9.43 | 6.16 | -62 | -20 | 28 |
| right supramarginal gyrus / tempoparietal junction | 557 | 7.24 | 5.31 | 60 | -26 | 28 |
| | | | IH2F1 | | | |
| left supramarginal gyrus / tempoparietal junction | 2633 | 6.18 | 4.8 | -62 | -30 | 36 |
| right supramarginal gyrus / tempoparietal junction | 1677 | 5.78 | 4.59 | 62 | -24 | 26 |
| right posterior middle temporal gyrus | 659 | 4.86 | 4.06 | 58 | -66 | -2 |
| left posterior middle temporal gyrus | 232 | 4.39 | 3.76 | 28 | 36 | -18 |
| right inferior frontal gyrus | 182 | 4.26 | 3.67 | 62 | 8 | 12 |
| left temporal pole | 12 | 3.93 | 3.45 | -56 | 12 | -10 |
| right inferior frontal gyrus | 16 | 3.77 | 3.33 | -60 | 4 | 12 |

**Table S1. Results of univariate fMRI analysis** (in association with **Fig. 2A.**) **A.** Neural activity during choices modulated by the Euclidean distance of inference trajectories via the hub ($E_{H2F1}$). **B.** Neural activity during choices modulated by the rank difference in the task-relevant dimension ($D_{H2F1}$) and the rank difference in the task-irrelevant dimension ($I_{H2F1}$) from the hub (1-D distance of inferences). All reported effects use threshold-free cluster enhancement (TFCE) corrected at $p_{TFCE}<0.05$.

| | Route through H2 | | | |
|---|---|---|---|---|
| | EH2F1 | DH2F1 | IH2F1 | AH2F1 |
| left Entorhinal Cortex | **0.82** ⋄ | 0.03 | 0.01 | 0.01 |
| right Entorhinal Cortex | **0.91** ⋄ | 0.00 | 0.00 | 0.00 |
| left ventromedial prefrontal cortex / medial orbitofrontal cortex | **0.89** ⋄ | 0.02 | 0.00 | 0.01 |
| right ventromedial prefrontal cortex / medial orbitofrontal cortex | **0.85** ⋄ | 0.03 | 0.00 | 0.01 |

| | Route through H1 | | | |
|---|---|---|---|---|
| | EH2F1 | DH2F1 | IH2F1 | AH2F1 |
| left Entorhinal Cortex | 0.02 | 0.02 | 0.00 | 0.04 |
| right Entorhinal Cortex | 0.00 | 0.00 | 0.00 | 0.08 |
| left ventromedial prefrontal cortex / medial orbitofrontal cortex | 0.01 | 0.01 | 0.00 | 0.03 |
| right ventromedial prefrontal cortex / medial orbitofrontal cortex | 0.01 | 0.03 | 0.00 | 0.03 |

| | Direct Route from F1 to F2 | | | |
|---|---|---|---|---|
| | EH2F1 | DH2F1 | IH2F1 | AH2F1 |
| left Entorhinal Cortex | 0.02 | 0.01 | 0.00 | 0.02 |
| right Entorhinal Cortex | 0.00 | 0.00 | 0.00 | 0.00 |
| left ventromedial prefrontal cortex / medial orbitofrontal cortex | 0.00 | 0.01 | 0.00 | 0.01 |
| right ventromedial prefrontal cortex / medial orbitofrontal cortex | 0.01 | 0.01 | 0.00 | 0.01 |

**Table S2. Exceedance probability (XP) computed from Bayesian model selection** (in association with **Fig. 2B.**) ⋄ indicates the winning model, indicating that $E_{H2F1}$ explains the variance of the activity in the ROI better than other different distance measures.

**A.**

| ROIs | Euclidean (E) | 1-D Relevant rank diff. (D) | 1-D Irrelevant rank diff. (I) | Context (C) | Group (G) |
|---|---|---|---|---|---|
| HC left | 0.091±0.007 *** | 0.078±0.005 *** | 0.051±0.006*** | -0.002±0.004 | 0.018±0.007 |
| HC right | 0.095±0.007 *** | 0.081±0.004 *** | 0.048±0.005*** | 0.004±0.003 | 0.012±0.008 |
| EC left | 0.066±0.009 *** | 0.064±0.005 *** | 0.031±0.005*** | 0.001±0.003 | 0.010±0.009 |
| EC right | 0.059±0.007 *** | 0.057±0.005 *** | 0.033±0.004*** | -0.006±0.003 | 0.011±0.007 |
| vmPFC/mOFC left | 0.064±0.007 *** | 0.048±0.008 *** | 0.049±0.009* | 0.015±0.008 | -0.001±0.005 |
| vmPFC/mOFC right | 0.067±0.008 *** | 0.046±0.009 *** | 0.044±0.009* | 0.014±0.007 | -0.006±0.006 |
| Amygdala left | 0.003±0.008 | 0.008±0.009 | 0.010±0.007 | -0.002±0.008 | 0.009±0.008 |
| Amygdala right | 0.008±0.006 | 0.007±0.007 | 0.019±0.008 | -0.004±0.006 | 0.002±0.007 |
| Motor left | -0.008±0.012 | -0.013±0.007 | 0.002±0.011 | 0.002±0.006 | 0.009±0.007 |
| Motor right | 0.009±0.010 | 0.003±0.007 | 0.008±0.007 | -0.003±0.006 | -0.008±0.007 |

**B.**

| ROIs | Within-Group E | Between-group (Hub) E | Between-group (NonHub) E |
|---|---|---|---|
| HC left | 0.182±0.010 *** | 0.237±0.010 *** | 0.053±0.009 *** |
| HC right | 0.188±0.012 *** | 0.235±0.010 *** | 0.051±0.009 *** |
| EC left | 0.160±0.011 *** | 0.241±0.012 *** | 0.031±0.011 |
| EC right | 0.149±0.012 *** | 0.253±0.012 *** | 0.024±0.011 |
| vmPFC/mOFC left | 0.066±0.011 *** | 0.173±0.018 *** | 0.000±0.014 |
| vmPFC/mOFC right | 0.067±0.011 *** | 0.184±0.019 *** | -0.001±0.012 |

**C.**

| ROIs | Within-Group E *vs.* Between-group NonHub E | Between-group Hub E *vs.* NonHub E |
|---|---|---|
| HC left | 4.54 *** | 4.54 *** |
| HC right | 4.54 *** | 4.54 *** |
| EC left | 4.52 *** | 4.54 *** |
| EC right | 4.37 *** | 4.54 *** |
| vmPFC/mOFC left | 3.41 *** | 4.37 *** |
| vmPFC/mOFC right | 3.39 *** | 4.54 *** |

**Table S3. Representational similarity analysis (RSA) in the region of interests (ROIs). A.** In association with **Fig. 4C-F**. the mean rank correlation (Kendall's $\tau_A$ ± s.e.m), which indicates the relatedness of the representational dissimilarity matrix (RDM) estimated in each ROI to the model RDM (E, D, I, C, and G) **B.** In association with **Fig. 4I**. The effect of E (rank correlation, Kendall's $\tau_A$ ± s.e.m) was separately estimated for the within-group pairs, the between-group pairs of hubs (a hub and the faces that were directly paired with the hub), and the between-group pairs of non-hubs. **C.** In association with **Fig. 4I**. The z-values computed from two-sided Wilcoxon signed-rank test which shows that the effect of E was stronger for within-group pairs and between-group pairs of hubs compared to the effect of E for between-group pairs of non-hubs in the ROIs. All FWE corrected with Bonferroni-Holm method for multiple comparisons, *** $p_{FWE}<0.001$, ** $p_{FWE}<0.01$, * $p_{FWE}<0.05$.

**A.**

| | Cluster size | T | Z | Peak coordinate (MNI) x | y | z |
|---|---|---|---|---|---|---|
| Model RDM | Pairwise Euclidean distance (E) | | | | | |
| right central/medial orbitofrontal cortex | 644 | 5.00 | 4.15 | 12 | 42 | -20 |
| right subgenural area | 481 | 4.97 | 4.13 | 12 | 14 | -20 |
| right entorhinal cortex | 32 | 4.78 | 4.01 | 20 | 0 | -36 |
| left lateral orbitofrontal cortex | 917 | 4.41 | 3.77 | -24 | 26 | -18 |
| right hippocampus | 193 | 4.37 | 3.75 | 30 | -6 | -18 |
| posterior cingulate cortex | 208 | 3.88 | 3.42 | -4 | -44 | 30 |
| posterior/medial cingulate cortex | 315 | 3.76 | 3.33 | 2 | -26 | 38 |
| right lateral orbitofrontal cortex | 127 | 3.09 | 2.82 | 28 | 24 | -18 |
| right visual cortex | 717 | 3.02 | 2.77 | 22 | -78 | 10 |

**B.**

| Model RDM | Pairwise rank difference in the task-relevant hierarchy (D) | | | | | |
|---|---|---|---|---|---|---|
| bilateral medial prefrontal cortex / subgenual area | 483 | 5.32 | 4.34 | -12 | 18 | -10 |
| bilateral posterior cingulate cortex | 578 | 5.27 | 4.31 | -4 | -40 | 30 |
| bilateral dorsomedial prefrontal cortex | 4483 | 5.09 | 4.20 | -10 | 52 | 10 |
| bilateral precuneus | 734 | 4.72 | 3.97 | 10 | -52 | 6 |
| left temporoparietal junction | 38 | 4.64 | 3.92 | -56 | -36 | 40 |
| left lateral orbitofrontal cortex | 1365 | 4.57 | 3.88 | -38 | 16 | -10 |
| right inferior frontal gyrus | 45 | 4.43 | 3.79 | 62 | 18 | 14 |
| right lateral orbitofrontal cortex | 231 | 4.28 | 3.69 | 28 | 22 | -18 |
| right dorsolateral prefrontal cortex | 107 | 4.24 | 3.66 | 24 | 64 | 20 |

**C.**

| Model RDM | Partialling out D from the RDM for E (E') | | | | | |
|---|---|---|---|---|---|---|
| bilateral posterior cingulate cortex | 13508 | 4.46 | 3.81 | -16 | -32 | 50 |
| right central/medial orbitofrontal cortex | 348 | 4.27 | 3.68 | 10 | 42 | -18 |
| dorsomedial prefrontal cortex | 82 | 4.10 | 3.57 | 8 | 58 | 36 |
| left precuneus | 832 | 3.77 | 3.34 | -24 | -18 | 58 |

| | | | | | | |
|---|---|---|---|---|---|---|
| left fusiform gyrus | 154 | 3.46 | 3.11 | -32 | -44 | -16 |
| left visual cortex | 199 | 3.26 | 2.96 | -42 | -86 | 18 |
| right hippocampus | 26 | 2.8 | 2.6 | 38 | -24 | 2 |

**Table S4. Whole-brain searchlight representational similarity analysis (RSA). A.** Brain regions in which the representational dissimilarity matrix (RDM) estimated by the searchlight analysis was predicted by the model RDM of pairwise Euclidean distances on the 2-D social space (E), (in association with **Fig. 4G**). **B.** Regions predicted by the model RDM of pairwise differences in the rank in the task-relevant dimension (D), (in association with **Fig. S8A**). **C.** Regions predicted by the model RDM of E' (in association with **Fig. 4H**). E' indicates E after partialing out confounding covariance with D (**Fig. S8B**).
